# The Ecosystem Integrity Index: a novel measure of terrestrial ecosystem integrity

**DOI:** 10.1101/2022.08.21.504707

**Authors:** S.L.L. Hill, M.L.K Harrison, C. Maney, J. Fajardo, M. Harris, N. Ash, J. Bedford, F.S. Danks, D. Guaras, J. Hughes, M. Jones, T. Mason, N. Burgess

## Abstract

While the importance of ecosystem integrity has long been recognised (Leopold, 1949), conservation science has tended to focus on measuring and monitoring species and habitats, avoiding the complexities of working at the ecosystem level. Ecosystems are highly dynamic, defined by both living and non-living components as well as their interactions (CBD, 1992), making it difficult to assess baseline levels of integrity. We present a novel index that represents the integrity of all terrestrial ecosystems globally at 1km^2^ resolution: the Ecosystem Integrity Index (EII). The index provides a simple, yet scientifically robust, way of measuring, monitoring and reporting on ecosystem integrity. It is formed of three components; structural, compositional and functional integrity, and measured against a natural (current potential) baseline on a scale of 0 to 1. We find that ecosystem integrity is severely impacted in terrestrial areas across the globe with approximately one fifth of all ecosystems and one quarter of all ecoregions having lost, on average, over half of their ecosystem integrity. At a national scale, we estimate similar challenges with 115 nations or territories having lost, on average, over half of their ecosystem integrity. This presents a significant threat for humanity as such levels of degradation are likely to be linked to substantial declines in the ecosystem services on which humanity is reliant. The EII has been developed principally to help national governments measure and report on Goal A of the Kunming-Montreal Global Biodiversity Framework (GBF) (CBD, 2022a), for which it has been listed as a Component Indicator. The EII will also be useful in helping non-state actors measure and report their contributions to the GBF and is listed as an indicator by both the Taskforce for Nature-Related Financial Disclosures (TNFD) (TNFD, 2023) and the Science Based Targets Network (SBTN) (SBTN, 2023). The EII aims to enable these actors to make informed decisions on the conservation, restoration and sustainable use of ecosystems for which they are wholly or partly responsible. We propose that with sufficient effort, ecosystem integrity can be restored and contribute towards the GBF’s vision of living in harmony with nature, resulting in the safeguarding of the ecosystem services on which humanity depends.

## Introduction

Ecosystem integrity encompasses the full complexity of an ecosystem, including the physical, biological and functional components, together with their interactions, and measures these against a ‘natural’ (i.e., current potential) reference level (Carter *et al*., 2019). Ecosystem integrity is fundamental to the stability of Earth systems on which humanity depends. For instance, natural areas containing ecosystems with higher integrity have greater potential to provide services such as carbon sequestration (Lewis *et al*., 2009), maintenance of water quality (Mello *et al*., 2018), climate regulation (Bonan, 2008), pest control (Bianchi *et al*., 2006), and pollination (Carvalheiro *et al*., 2010) – as well as supporting higher levels of biodiversity (Gibson, 2011; Barlow *et al*., 2016).

The importance of healthy ecosystems (ecosystems with high integrity) is recognised in the three Rio Conventions: the Convention on Biological Diversity (CBD), the UN Framework Convention on Climate Change and the UN Convention to Combat Desertification.

The Kunming-Montreal Global Biodiversity Framework (GBF), negotiated under the CBD, includes ambitions for the maintenance, restoration or enhancement of ecosystem integrity within Goal A (CBD, 2022a). This index has been designed to help parties to the Convention to directly monitor ecosystem integrity and is listed as a Component Indicator within the adopted Monitoring Framework for the GBF (CBD, 2022b).

Meeting the goals and targets of the GBF will require action not only at the national level (by the 196 parties to the CBD), but also by a range of other actors including businesses. The Ecosystem Integrity Index (EII) provides a means of alignment between the goals and targets of the GBF and also; 1) target guidance under development by the Science Based Targets Network (SBTN); 2) the risk management and disclosure framework of the Taskforce on Nature-related Financial Disclosures (TNFD). Both TNFD and SBTN have identified the need for a scalable metric of ecosystem integrity, linked to the GBF, which is readily usable by businesses and financial institutions for evidence-based target setting, monitoring and disclosure (TNFD, 2022; SBTN, 2020).

### Conceptualisation of ecosystem integrity

The notion of ecosystem integrity first emerged in the work of Leopold (1949), who used the term to encompass the basic criteria necessary for biotic communities to remain stable. The concept was elaborated upon by Parrish *et al*. (2003), who proposed that ‘an ecological system has integrity when its dominant ecological characteristics (e.g., elements of structure, composition, function, and ecological processes) occur within their natural ranges of variation’. Carter *et al*. (2019), simplified this to define ecosystem integrity as ‘the extent to which the composition, structure, and function of an ecosystem fall within their natural range of variation’. Structural integrity comprises the three-dimensional aspect of ecosystems – the biotic and abiotic elements that form the heterogeneous matrix supporting composition and functioning. Composition refers to the biotic constitution of ecosystems – the pattern of the makeup of species communities. Functioning describes the ecological processes and ecosystem services provided by the ecosystem. It follows that any measurement of ecosystem integrity should encompass structural, compositional, and functional components. However, it should be noted that the three components are interdependent and are likely to covary with varying pressures on the system.

Ecosystem integrity has generated considerable interest in recent years with the development of several indices designed to assess changes in ecological factors relating to integrity at a global level or within specific habitat types. Recent work has focussed on measuring ecosystem extent (there remains little scientific consensus on the best way to do this (Sayre *et al*., 2020; IUCN-CEM, 2022)), and on aspects of ecosystem integrity (Beyer *et al*., 2019; Blumetto *et al*., 2019; Grantham *et al*., 2020; Hansen *et al*., 2021; Mora, 2017; Perkl, 2017). For instance, Grantham et al. (2020) created the Forest Landscape Integrity Index (FLII) integrating data on forest extent, observed and inferred human pressures, and changes in forest connectivity, and an index of ecosystem integrity – focussing on agricultural systems – was developed by Blumetto *et al*. (2019) to monitor integrity at the farm level. (For a longer discussion of relevant work in this area, see SI – methodology).

Despite these various efforts, there is currently no index that matches the definition of ecosystem integrity, in that it brings together all three components of ecosystem integrity, being structure, composition and functioning, measured against a natural baseline, that can be applied globally.

### Introducing the Ecosystem Integrity Index

Our Ecosystem Integrity Index (EII) quantifies degradation to ecosystem integrity based on an aggregation of all three components: ecosystem structure, composition, and functioning. This framing is consistent with the definitions provided by Carter *et al*. (2019) and Parrish *et al*. (2003). Our index provides a measure of a fundamental property of ecosystems, ecosystem integrity, at a fine scale, 1km^2^, but can be aggregated to other levels, increasing its utility to provide support to monitoring and spatial planning at local to global scales. The ability to aggregate EII at different scales allows it to be applied in multiple use-cases. For example, it is compatible with all levels of the IUCN Global Ecosystem Typology (Keith *et al*., 2020), which forms the basis for the biomes used within the TNFD Recommended Disclosures (TNFD, 2023), or, as we demonstrate in this analysis, it can be used with the World Ecosystems typology (Sayre *et al*., *2020)* or ecoregions (Dinerstein *et al*., 2017), but it is also compatible with national-level ecosystem typologies and so could support governments engaged in spatial planning.

The first component of EII – structure – is correlated with habitat intactness, the area of contiguous habitat, and fragmentation. This component is calculated by inputting the Human Modification Index (HMI) (Kennedy *et al*., 2019) into the algorithm described in Beyer *et al*. (2019) that accounts for habitat intactness within a grid cell as well as the influence of habitat quality within neighboring cells. This method ensures that our final structure layer encompasses impacts of landscape fragmentation as well as local intactness.

The second component captures ecosystem composition, which refers to the identity and variety of life within an area. The metric chosen for this layer is the Biodiversity Intactness Index (BII), which summarizes change in the make-up of local ecological communities in response to human pressures (Newbold *et al*., 2016, Hill *et al*., 2019). The BII is calculated using two models fitted to species community data recorded in the PREDICTS database (Hudson *et al*., 2017). The first model estimates the impact of human pressures on the total abundance of species within a community and the second analyses the similarity between the relative abundance of each of the species in degraded areas against those of the same communities found within natural areas. The product of the two models, projected onto maps of human pressures, results in the BII.

The third component captures ecosystem function. The functioning component is estimated using the difference between potential natural and current net primary productivity (NPP) within each 1km^2^ grid cell. Current NPP values are taken from remote sensed geospatial layers (Running and Zhao, 2019). Potential natural NPP values are modelled using environmental input data (for model details see Methods).

The three component layers are aggregated to form the EII. We used a modified minimum value approach, in which we take the lowest score of the structure, composition and functioning components as the EII value for a pixel and then down-weight the value based upon the integrity of the other two components. This method was chosen with the reasoning that the integrity of an ecosystem cannot be higher than the minimum score from any of the three components. The derived EII, as well as the three components, ranges between 0 and 1, with the most degraded areas having a score approaching zero.

## Results

Ecosystem integrity is highly degraded across much of the world with only remote areas in higher latitudes (note that Antarctica was excluded from the analysis), remote areas of rainforest across the tropics, and extremely arid areas across the planet estimated to have extensive areas where ecosystem integrity has been retained (Figure 1A).

**Figure 1.**
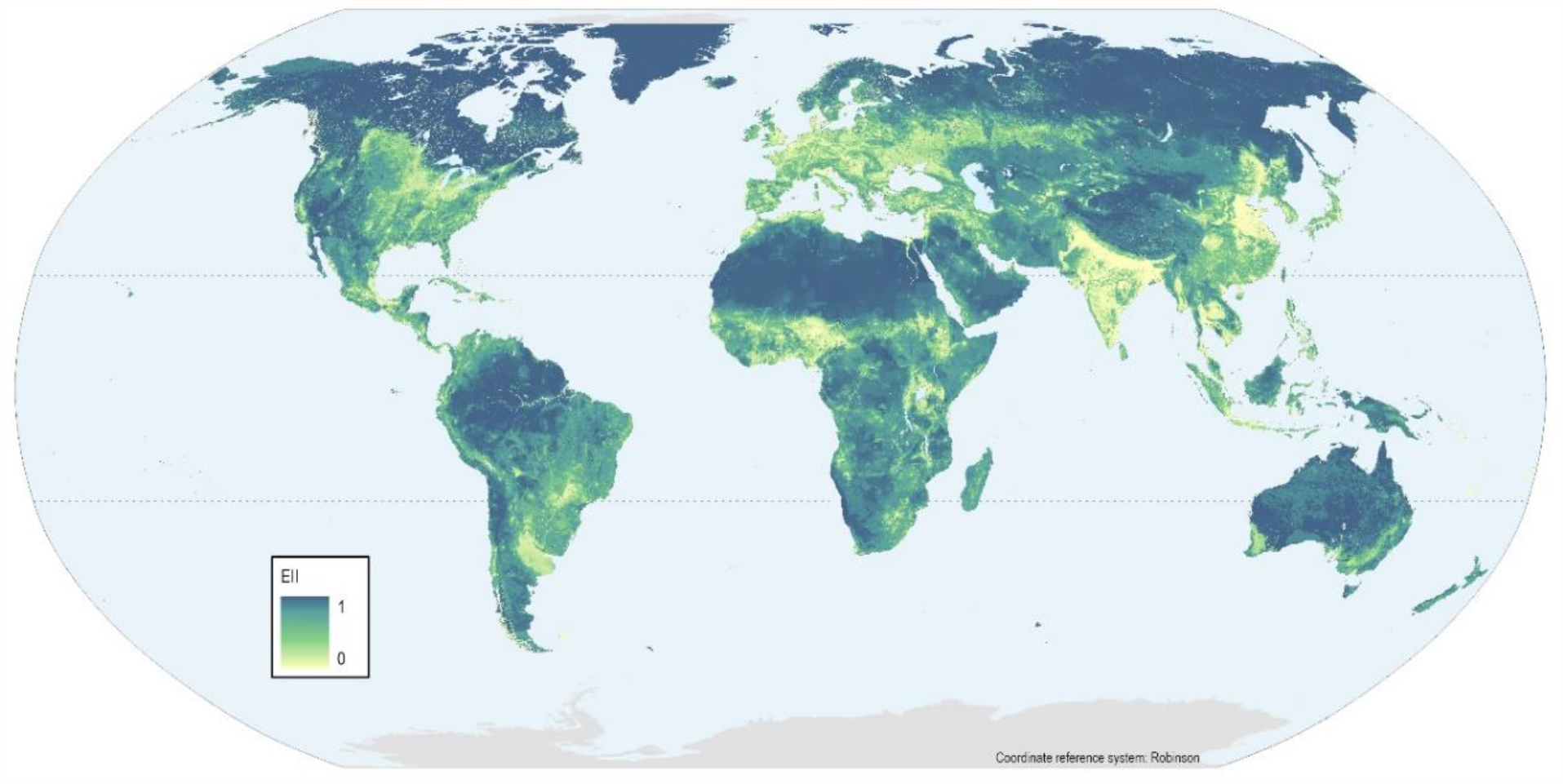
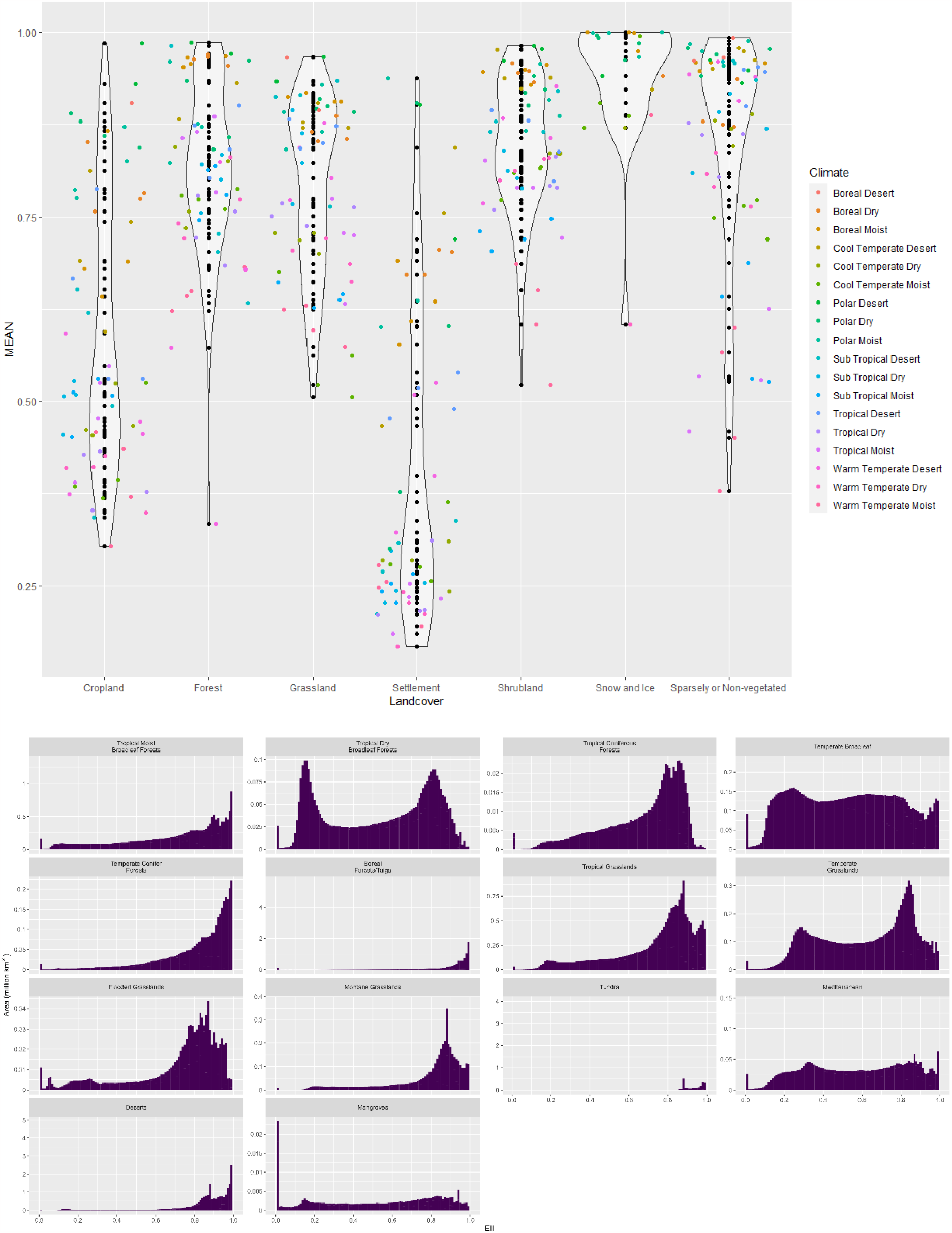
Global ecosystem integrity. A) The Ecosystem Integrity Index for all terrestrial areas. Areas with high levels of ecosystem integrity are shown in dark blue, whereas those with highly degraded ecosystem integrity are shown in pale yellow. Note: the boundaries and names shown and the designations used on this map do not imply official endorsement or acceptance by the United Nations. B) Mean ecosystem integrity per ecosystem (Sayre *et al*., 2020) grouped by landcover types. Colours indicate climate regions. C) Frequency of ecosystem integrity index values within terrestrial biomes (WWF 2017).

Mean ecosystem integrity across all ecosystems (ecosystem boundaries as defined by Sayre *et al*., 2020) was 0.73 (±0.22), with 74 of 422 ecosystems having, on average, lost at least half of their ecosystem integrity. Ecosystems within human settlement areas in warm temperate regions were found to have the lowest levels of ecosystem integrity, followed by ecosystems found within cropland areas (Figure 1B). At broader scales, approximately one quarter (198 of 824) of ecoregions have, on average, lost at least half of their ecosystem integrity (mean EII <0.5). Dry tropical broadleaf forests are more degraded than moist, and temperate grasslands, temperate broadleaf forest and Mediterranean forests, woodlands and scrub all have high levels of degradation (Figure 1C).

Ecosystem degradation is widespread at the national scale (Figure 2). Across all 255 countries and territories assessed, the mean EII value was 0.55 (SD:0.23). When comparing between continents, North American countries and territories were found to have, on average, the lowest EII scores with a mean value of 0.46 (SD:0.22), and those in West Asia had the highest EII scores with a mean value of 0.68 (SD:0.17).

**Figure 2.**
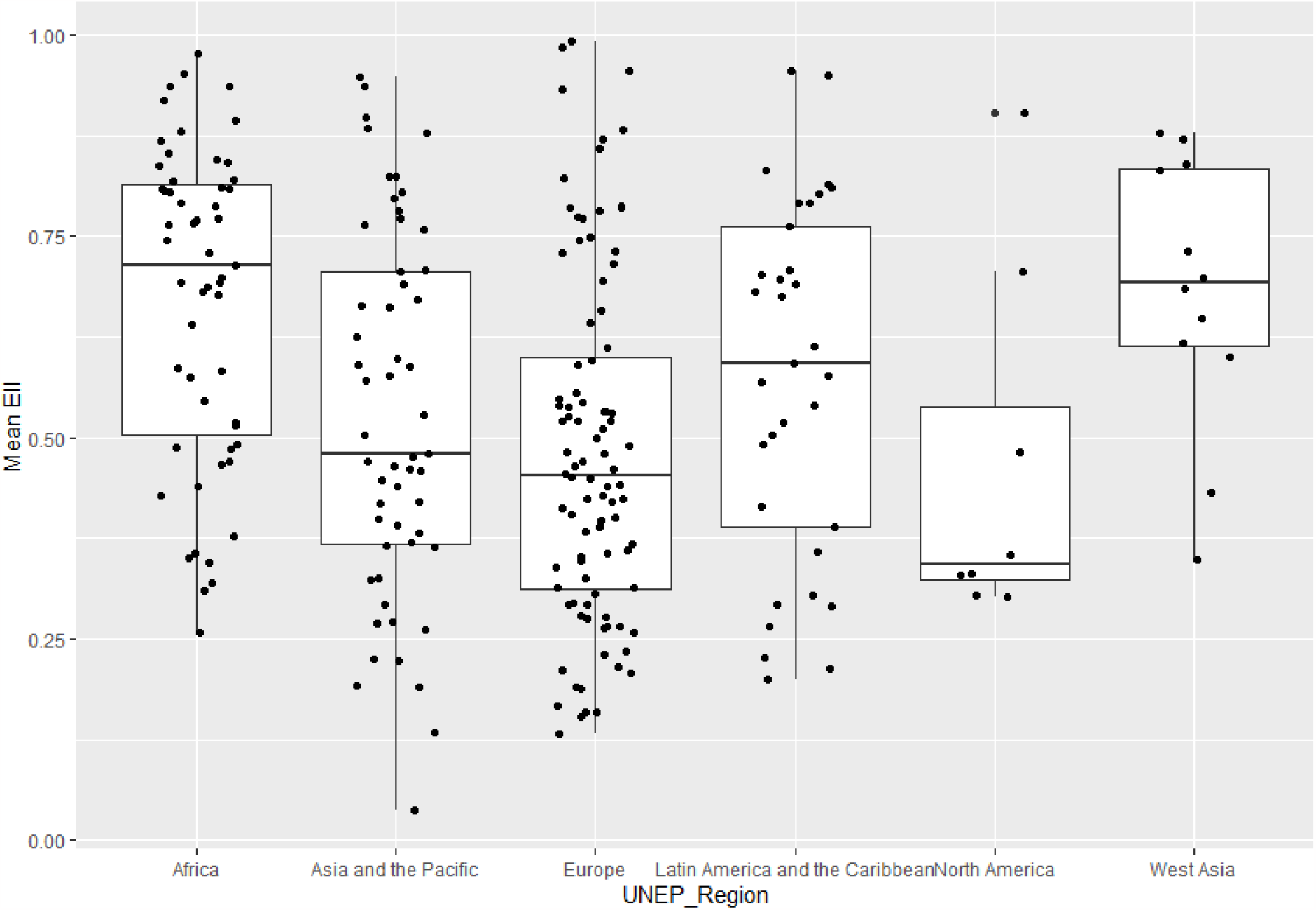
Ecosystem integrity within countries and territories grouped by UN Regions. Each dot represents the values for a single country. Box plots indicate mean ±SE.

## Discussion

The EII provides a unique insight into the extent that terrestrial ecosystems have been degraded across the world. We have used ecological theory and the best available data to create an index at operational scales (1km^2^) globally. Our results show that ecosystem integrity has been highly degraded across the terrestrial world with nearly one fifth of all ecosystems, and one quarter of all ecoregions, having lost, on average, over half of their ecosystem integrity. Such dramatic losses of ecosystem integrity are likely to have had severe consequences for species persistence and provision of ecosystem services, but further work is needed to understand how the spatial distribution of ecosystem integrity interacts with these factors. Such knowledge will be required to enable effective spatial planning, land management options and restoration interventions to help achieve the goals and targets of the Kunming-Montreal GBF.

We developed this index for two main use cases. The first is for use at the national and sub-national scales by countries seeking ways to monitor progress against their national goals and targets within the context of the Kunming-Montreal GBF. In particular, the index will facilitate tracking of progress towards national contributions to Goal A, maintenance, enhancement and restoration of ecosystem integrity, as well as targets 1, 2 and 10 covering spatial planning, restoration and sustainable use.

The global EII layer will be made available through assessment tools such as the UN Biodiversity Lab (https://unbiodiversitylab.org/) to support countries in planning for the delivery of these targets and monitoring progress in terms of ecological outcomes.

Our second use case is for businesses and financial institutions which are starting to include biodiversity measurement in their risk assessment and performance reporting. There is an urgent need for trackable targets using an appropriate set of metrics to enable the private sector to move towards positive outcomes for biodiversity (Locke *et al*., 2022; Milner-Gulland, 2022). Such outcomes are reliant upon the ability of a business to measure their biodiversity baseline, assess their exposure to biodiversity risk within operations, and to drive innovation towards positive outcomes on the ground through a quantified understanding of the pressure-response relationship. Initiatives such as the SBTN and the TNFD are working with businesses and financial institutions, helping them contribute to nature-positive outcomes. They are developing guidance for businesses to set science-based targets, and through recommendations for disclosures when reporting on impacts and dependencies on nature. In both these initiatives, participating businesses and financial institutions will need to understand where they interact with ecosystems and identify priority locations, for example, in the ‘Locate’ step in the TNFD LEAP (Locate, Evaluate, Assess, Prepare) approach (TNFD, 2023) and within the ‘Assess’ and ‘Interpret and Prioritize’ steps of the SBTN’s 5-step approach (SBTN, 2020). The TNFD, requires under ‘Strategy Disclosure D’ (TNFD, 2023) that the location of assets or activities are disclosed for ‘priority locations’, defined as, inter alia, those which have ‘high ecosystem integrity’ and those with ‘rapid decline in ecosystem integrity’. When implementing actions, and reporting on results, businesses and financial institutions will also need to understand their potential impacts and dependencies on ecosystems before going on to monitor and disclose how the integrity of these ecosystems is increasing or decreasing over time. The EII can support all of these use cases.

In the case of both state and non-state actor users, the EII should be seen as complementary to, and ideally used in conjunction with, measures of other facets of biodiversity. Complementary species metrics, for instance, include measures of species importance and threat across the world, such as the Species Threat Abatement and Restoration (STAR) metric based on the International Union for Conservation of Nature (IUCN) Red List (Mair *et al*., 2021), or range rarity metrics such as those presented in Hill et al. (2019). The Red List of Ecosystems (RLE) provides a framework for assessment and prioritisation of ecosystems analogous to the species threat assessments used to inform the Red List of Species (Rodriguez *et al*., 2015). The EII can be paired with the RLE to provide standardised information on ecosystem integrity to guide RLE assessments, or it can be used to complement the information provided by the RLE by providing spatially explicit information on ecosystem integrity across different ecosystems that is regularly updated.

The index has been designed to utilise datasets that track trend data, allowing annual updates as well as the potential to calculate historical trendlines. The EII framework also has the flexibility to allow exploration of different management practices at a local level by businesses, or pathways to sustainability with the alignment of the EII framework to national or global scenarios. The index can be disaggregated into its component layers and this may be informative when determining underlying causes of low ecosystem integrity; however, we encourage its use as an aggregated index, with actions focussed on improving integrity as a whole, rather than just one aspect of ecosystem health. Although many nature-positive actions will be beneficial for all three components of integrity, it should be recognised that components may respond at different rates in terms of recovery towards a more natural state. These temporal lags should be considered when planning actions and in the tracking of progress towards targets for nature.

The EII has limitations in that it does not yet directly quantify degradation to ecosystem integrity caused by climate change (although recent climate change impacts may contribute to lower scores through changes to NPP as reflected in the functioning layer). We note that many of the high latitude areas that appear unimpacted in this analysis may be at particular risk from climate change (Beaumont *et al*. 2011; Asamoah *et al*. 2021). Another limitation is that the composition and structural components are derived from models that assume the same pressure response relationships across all terrestrial ecosystems (as per the layers from which these components are derived). Further work will target these limitations to examine the sensitivity of the components to these assumptions.

Research currently in progress aims to 1) calculate EII for freshwater and marine habitats, 2) link EII with risk to business operations through loss of ecosystem services, and 3) assess business counterfactuals for EII. Future development of EII could be enhanced by linking the models used here to emerging temporal 10m land cover products (for instance, the Sentinel-2 10m Land Use/Land Cover Time Series; Karra *et al*., 2021 or the Dynamic World layer; Brown *et al*., 2022) or to ground-based biodiversity monitoring to provide higher resolution information on biodiversity change at the local scale, enabling more regular updates of the EII.

The EII shows dramatic degradation to ecosystem integrity over much of the terrestrial area globally. It has been designed to be flexible, responsive and updateable, and useful at multiple scales. As a metric it provides an important means for countries and businesses to measure ecosystem integrity within areas of interest, to set targets and to take action to enhance ecosystem integrity at appropriate geographical scales.

## Methods

### Structural integrity

An ideal measurement of structural integrity should capture both complexity and the multi-dimensionality of an ecosystem. This should include a measure of how intact the vertical or internal structure of a given location is, as well as the intactness of such features in the surrounding landscape (Carter *et al*., 2019; Grantham *et al*., 2020; Ehbrecht *et al*., 2021; Rayfield *et al*., 2011).

A single range of measurable characteristics is unsuitable for measuring ecosystem structural integrity at the global scale because the ‘natural’ state of ecosystem structure varies greatly by realm, ecoregion, and habitat type (e.g., Tierney *et al*., 2009). We therefore focus the structural integrity component on capturing the landscape level intactness through a knowledge of where human pressures are assumed to cumulatively degrade structural integrity (e.g., Venter *et al*., 2016;

Kennedy *et al*., 2019), combined with an adaptation of the methods established by Beyer *et al*. (2019) to produce a layer that is sensitive to habitat patch size, fragmentation, and connectivity. This is achieved by comparing each location on a map not just to a natural reference for that point, but the whole neighbourhood around that point to a counterfactual landscape in which no pressures occur, computed at a scale in agreement with other connectivity metrics (e.g., Saura *et al*., 2018).

To measure structural integrity, we used the Human Modification Index (HMI) layer (Kennedy *et al*., 2019) that is based upon the following pressure layers: croplands, pasturelands, rangelands, plantations, built-up areas, human population density, roads, rails, quarries and mining, wind turbines, electrical infrastructure and powerlines (SI). We used the same pressure intensities described in Kennedy *et al*. (2019). Second, we inverted the human modification layer to transform it in an equivalent of the habitat quality and extent layer in Beyer *et al*. (2019). Finally, we applied the Beyer *et al*. (2019) method to incorporate landscape quality and fragmentation through the use of a moving window that compares the quality of a grid cell with that of cells in their neighbourhood, which we defined as cells within a 27km distance (i.e., grid cells in a 55km x 55km box were used to estimate the structural integrity of the target cell at the centre of the box). All spatial processing was done in Google Earth Engine with a resolution of 30 arcsec.

Our layer differs from the one produced in the Beyer *et al*. (2019) paper due to the source ‘habitat quality’ layer. Where the original authors use the Human Footprint Index (HFI) (Venter *et al*., 2016), rescaled with an exponential function, as their layer, we opt for a recreation of the HMI as described by Kennedy *et al*. (2019). This approach has the benefit of meaningful interpretation: an area with a value of 0.5 has twice the ‘intactness’ of an area with a value of 0.25. It also avoids the double use of neighbourhood functions. For example, the HFI takes roads and applies a buffer to estimate the area of their impact; when the landscape-level effects are calculated this area is then further expanded, which may overestimate the spatial extent of these threats to ecosystem structure. In contrast, the HMI uses the physical footprint of these threats, preventing this issue when comparing at the landscape scale.

### Compositional integrity

A measure of compositional intactness should incorporate elements of both species’ abundance and community composition (Haase *et al*., 2018; Carter *et al*., 2019). The Biodiversity Intactness Index (BII) has been suggested as a suitable metric to track global planetary boundaries due to its relevance to ecosystem health and functioning (Steffen *et al*., 2015; Mace *et al*., 2018). The BII estimates the change in the makeup of species communities relative to an ‘unimpacted’ baseline (Newbold *et al*., 2016). The BII tracks changes in the relative abundance of species within a community (excluding novel species) due to land use changes as well as other human pressures. The BII is based upon modelled relationships estimated using a large taxonomically and geographically representative database, the PREDICTS database (Hudson *et al*., 2017). Data representing the communities of sites from many land-uses and many taxonomic groups are used to construct hierarchical models for the two elements of the BII, abundance and compositional similarity. These modelled relationships are then projected onto global pressure maps to form the BII layer.

Mixed effects models were fitted using land use and density of human population as fixed effects (for full modelling details see SI) and block nested within study as random effects. A novel land use map was constructed using 1) land use data from the Historic Land Dynamics Assessment+ (HILDA+) project (Winkler *et al*., 2021), 2) the Spatial Database of Planted Trees (Harris *et al*., 2019), 3) the Global Land Analysis and Discovery (GLAD) database (Potapov *et al*., 2022) and 4) the Potential Threats to Soil Biodiversity data from the Global Soil Biodiversity Atlas (JRC, 2016). The HILDA dataset provides land use details denoting natural areas, cropland, pasture/rangeland and urban from 1899 to present. We supplemented this data as follows: 1) within current day natural areas we tracked transitions and noted time since transition back to natural to distinguish primary vegetation, mature secondary vegetation, intermediate secondary vegetation and young secondary vegetation, 2) within agricultural areas we distinguish use intensity using levels of soil threat, 3) we distinguish between pasture and rangeland using overlap with the GLAD database, and 4) we distinguish plantation forests using the SDPT. (For further details on how the land use map was constructed see SI).

### Functional integrity

Attempts at mapping ecosystem functioning have explored changes in the values of specific ecosystem function variables. One example comes from the framework assessing Human Appropriation of Net Primary Productivity (HANPP), which explores departures from potential NPP values to investigate where human production systems have modified ecosystem functioning and the extent of this change (Mayer *et al*., 2021). Other authors have taken a very different approach to functioning through mapping patterns in the diversity of functional traits. For instance, Faurby and Svenning (2015) estimated changes to the richness of mammalian functional traits. However, no single metric has yet been developed that is considered to represent all aspects of ecosystem functioning.

In the absence of current alternative more comprehensive methods, we opted for adapting the approach to focus on one key ecosystem function, NPP, because of its well-documented association with ecosystem functioning (Malhi *et al*., 2011; Mayer *et al*., 2021), its advantages in terms of spatial resolution (layers are available in raster format at 1km^2^ resolution or higher) and update frequency (e.g., MODIS releases a global new NPP layer annually). Indeed, within the planetary boundaries framework, HANPP, as a fraction of Holocene levels of NPP, is the metric of choice to measure functional biosphere integrity (Richardson *et al*., 2023). NPP is also known to be closely associated with the provision of key ecosystem services such as mass flow regulation and atmospheric regulation, pest control and water purification (Harrison *et al*., 2014).

We developed a global layer of potential NPP used as a ‘natural’ reference and compared it with a current-day NPP layer derived from remote sensing to map changes in NPP. Natural levels of NPP were modelled per grid cell using environmental variables trained on mean NPP levels (Running and Zhao, 2015) measured within strictly managed protected areas (IUCN categories I and II) between 2001 and 2021 (UNEP-WCMC and IUCN, 2017). We used a generalised linear model framework, with a Gamma distribution for the response variable and a log-link. Variables were selected for testing based on a literature review of likely predictors of NPP and included: latitude, bioclimatic variables (Brun et al. 2022); mean, min and max solar radiation (Fick and Hijmans, 2017); aridity (Global Aridity Index, Zomer and Trabucco, 2022); total nitrogen, cation exchange capacity, predicted sand concentration, pH of water in soil (Poggio *et al*., 2021); continuous heat-insolation load index (CHILI, Theobald *et al*., 2015); roughness of terrain, slope, topographic position index, terrain ruggedness index (Amatulli *et al*., 2018); and landforms (Sayre *et al*., 2020). As the selection process did not result in a single top model, spatial projections of the two best models were used as inputs to the NPP calculation, along with two previously published potential NPP layers (Brun *et al*. 2022; Mayer *et al*. 2021).

As the four potential NPP models provide a range of values, we included two types of uncertainty in our calculations. Firstly, for each grid cell, we calculated the range of possible values using the lowest value and highest value observed between all models. All measures were calculated as movements away from this range (i.e., where NPP was estimated to be lost, the value was compared to the lowest potential value, and where NPP was estimated to be gained, the value was compared to the highest potential value). Secondly, we accounted for inter-seasonal variation by modelling the coefficient of variation of annual measures between 2001 and 2021 following the same methodology as that undertaken for the potential NPP. This provided an estimation of what natural level of variation may be expected within each grid cell and allowed us to adjust our calculations to account for movements outside the natural levels of variation.

Ecosystem functioning occurs on different scales. To assess changes to ecosystem functioning on a broader scale the absolute loss or gain of NPP per grid cell was calculated; however, to account for changes to ecosystem functioning on a local scale the proportion of NPP measured versus potential per grid cell was calculated. The mean of the proportional change and the decile value of the absolute change provided the final functional integrity score.

### Aggregation to derive the global Ecosystem Integrity Index

To calculate the Ecosystem Integrity Index (EII) value per grid cell we first extracted the lowest scoring of the three ecosystem components within that grid cell. This calculation follows the logic that all components of integrity are integral and therefore the overall condition of the ecosystem is limited by the lowest component. This emphasizes the interconnected nature of ecosystem integrity and avoids issues that may have been caused by an averaging/additive approach. For example, phenomena such as extinction debt (Kuussaari *et al*., 2009) suggest it would be prudent to refrain from allowing moderate values where any one component has been significantly weakened.

However, simply taking the lowest value cannot distinguish between an area where all three components are degraded versus an area where only a single component is degraded. We therefore adjusted the EII value using fuzzy algebraic summing (Zadeh, 1965, see SI) to downgrade the EII score based on the level of degradation of the two components that were not providing the lowest score.

Mean EII scores were calculated at biome and ecoregion levels using terrestrial biomes and ecoregions spatial data (Dinerstein *et al*., 2019), and ecosystem levels using Sayre *et al*., (2020).

## Acknowledgements

We would like to thank Mike Harfoot, Sharon Brooks, Sebastian Bekker, Maria Julia Oliva, Tom Evans, Tom Brooks, Emily Nicholson, David Keith, Malcolm Starkey, Craig Beatty, Marco Daldos Pirri, Varsha Vijay and Pamela Collins for their comments and review of the EII methodology. This output has been funded in whole or part by the UK Research and Innovation’s Global Challenges Research Fund under the Trade, Development and the Environment Hub project (project number ES/S008160/1) and by the Science Based Targets Network.

