## Supplementary material for "The Ecosystem Integrity Index: a novel measure of terrestrial ecosystem integrity": SI

**Online Annex**

**Supplementary information – further methodology**

**Table S1**: data layers used in EII

| Pressure / layer class | Dataset provider | Year | Source |
|---|---|---|---|
| **Structure** | | | |
| **Global human modification of terrestrial systems** | SEDAC | 2016 (median year) | https://sedac.ciesin.columbia.edu/data/set/lulc-human-modification-terrestrial-systems/data-download |
| **Composition** | | | |
| **HILDA+** | PANGAEA | 2019 | https://doi.pangaea.de/10.1594/PANGAEA.921846 |
| **Spatial Database of Planted Trees (SDPT)** | WRI | 2019 | https://data.globalforestwatch.org/documents/gfw::planted-forests/about |
| **Human population density** | SEDAC | 2015 | https://sedac.ciesin.columbia.edu/data/collection/gpw-v4 |
| **PREDICTS database** | NHM | NA | https://data.nhm.ac.uk/dataset/?tags=predicts |
| **Soil Biodiversity Threats map** | ESDAC | 2016 | https://esdac.jrc.ec.europa.eu/content/global-soil-biodiversity-maps-0 |
| **Cropland expansion** | GLAD | 2019 | https://glad.umd.edu/dataset/croplands |
| **Functioning** | | | |
| **NPP** | NASA LP DAAC at the USGS EROS Center | 2021 | https://lpdaac.usgs.gov/products/mod17a3hgfv061/ |
| **Mean annual temperature (bio1)** | Chelsa | NA | https://chelsa-climate.org/exchelsa-extended-bioclim/ |
| **Mean temperature of the coldest quarter (bio11)** | Chelsa | NA | https://chelsa-climate.org/exchelsa-extended-bioclim/ |
| **Annual precipitation (bio12)** | Chelsa | NA | https://chelsa-climate.org/exchelsa-extended-bioclim/ |
| **Nitrogen** | Soilgrids | NA | https://soilgrids.org/ |

| | | | |
|---|---|---|---|
| **Biome** | WWF | NA | https://www.worldwildlife.org/publications/terrestrial-ecoregions-of-the-world |
| **Sand** | Soilgrids | NA | https://soilgrids.org/ |

<u>Additional ecosystem integrity work used to frame our approach</u>

Within different habitats and ecological interactions, a number of studies have developed ecological integrity indexes. As examples:

Grantham *et al*. (2020) created the Forest Landscape Integrity Index (FLII) integrating data on forest extent, observed and inferred human pressures, and changes in forest connectivity. Using FLII, they found that only 40% of the world's forests are of high integrity. However, this measure remains limited to forest ecosystems and does not consider composition or functioning. Similarly, Perkl (2017), developed an approach that uses fuzzy logic to model patterns of anthropogenic impact at large-scale landscape levels as well as at finer scales. This measure of landscape integrity, like the FLII, focuses primarily on the impact of infrastructure to quantify the level of human modification but does not account for changes in composition or functioning (Perkl, 2017).

Mora (2017) developed an approach for modelling ecological integrity, using spatial data layers and structural equation modelling, with a focus on measuring the extent to which ecological interactions were maintained in predator-prey systems. This method used a range of layers that provided information on diversity, as well as those that characterised habitat and connectivity, but none that explicitly measured ecosystem functioning (Mora, 2017).

An index of ecosystem integrity for agricultural systems was developed by Blumetto *et al.* (2019) to monitor integrity at the farm level. This semi-quantitative approach involves the visual assessment of plant species, vegetation structure, soil and riparian habitat, with each of these variables valued on a scale of 0–5, with 5 signifying the best condition (Blumetto *et al.,* 2019). Measuring some aspects of structure, composition and functioning, Blumetto et al.'s approach represents a simplified implementation of the concept of ecosystem integrity, but with a focus on only one system and at the local scale.

<u>Creating the BII – detailed methods</u>

Species abundance and composition similarity were built from the PREDICTS database (Hudson *et al.* 2017) as outlined in Hill *et al.* (2018). The data used included 3.85 million rows and information on approximately 31 000 taxa from 32 000 sites and 767 studies. Land use and land use intensity were taken from the original source publications, as registered in the database. Human population density was extracted for each PREDICTS site from GRUMP (CIESIN 2011).

We calculated the total species abundance as the summed number of individuals of all species for each site, and composition similarity using the asymmetric Jaccard index (Chao *et al.* 2004). This metric used the abundance in sites classified as having primary vegetation with 0 human population density as a baseline reference.

Local abundance and compositional similarity models were built using a mixed-effects modelling framework with the R package lme4 (Bates *et al.,* 2017). Abundance was square root transformed. For all models, site information relative to source, study and block information were used as random intercepts, as selected using Akaike's Information Criterion (AIC) (Zuur *et al.*, 2009). The abundance model selection process included human population density (log-transformed), land use, land-use intensity and the interaction of human population density and land use as fixed effects. Fixed effects were selected using backwards stepwise selection based on AIC values (Zuur *et al*., 2009). The compositional similarity model replicated the framework detailed in De Palma (2018). It used pairwise comparisons between all sites based on four variables: Jaccard's asymmetric compositional similarity (logit transformed), geographic distance (log transformed), environmental distance, and difference in human population density (log transformed). Contrasts including the intact baseline were used to build the model. Compositional similarity was modelled against the other three variables.

Both models were projected using raster data for predictors at 1km$^2$ resolution to output gridded maps of species abundance and composition similarity. BII is obtained from the multiplication of both projected models (Newbold *et al*., 2016).

<u>Aggregating EII – detailed methods on fuzzy algebraic sums</u>

Fuzzy algebra provides a means of accounting for complex and uncertain relationships between members of sets (Zadeh 1965) and is often used with spatial data to allow more flexible combinations of GIS information (Pradhan *et al.,* 2009). When aggregating EII values across the three components, we used the formulas:

1. "$x_i = MIN(a_i, b_i, c_i)$"
2. "$EII = (x_i * (y_i * z_i))$

**Supplementary information - results**

**Table S2** shows the full ranked list of countries and territories in terms of their EII values. As can be seen many of the countries and territories with the lowest EII values are city states, small islands, and other countries and territories with relatively low surface and relatively large rural populations.

| Country or territory | Mean EII |
|---|---|
| Nauru | 0.03661 |
| Bermuda | 0.13123 |
| Niue | 0.134519 |
| San Marino | 0.154019 |
| Jersey | 0.157958 |
| Guernsey | 0.159463 |
| British Virgin Islands | 0.167193 |
| New Caledonia | 0.187491 |
| Saint Pierre et Miquelon | 0.190313 |
| Vanuatu | 0.190534 |
| Bangladesh | 0.191723 |
| Saint Vincent and the Grenadines | 0.199391 |
| Falkland Islands (Malvinas) | 0.207265 |
| Réunion | 0.210948 |
| Saint Kitts and Nevis | 0.213354 |
| Faroe Islands | 0.215674 |

| | |
|---|---|
| Tonga | 0.222784 |
| Singapore | 0.225501 |
| Grenada | 0.226448 |
| Moldova | 0.230999 |
| Turks and Caicos Islands | 0.235175 |
| Montserrat | 0.257537 |
| Comoros | 0.258083 |
| Cook Islands | 0.26184 |
| Saint Martin | 0.26384 |
| Pitcairn | 0.265181 |
| Haiti | 0.265394 |
| Martinique | 0.265576 |
| Norfolk Island | 0.269551 |
| Christmas Island | 0.270934 |
| Denmark | 0.274931 |
| Bonaire | 0.277214 |
| Belgium | 0.278526 |
| Dominica | 0.290576 |
| Luxembourg | 0.292519 |
| India | 0.292638 |
| Antigua and Barbuda | 0.29309 |
| Mayotte | 0.293145 |
| Guadeloupe | 0.294636 |
| United States Virgin Islands | 0.302006 |
| El Salvador | 0.303867 |
| Guam | 0.304375 |
| Hungary | 0.306102 |
| Burundi | 0.309498 |
| Anguilla | 0.312619 |
| Liechtenstein | 0.314291 |
| Chagos Archipelago | 0.319957 |
| Solomon Islands | 0.323889 |
| Samoa | 0.324657 |
| Netherlands | 0.324784 |
| Puerto Rico (USA) | 0.329493 |
| American Samoa | 0.331055 |
| Saint Barthélemy | 0.338498 |
| Nigeria | 0.344455 |
| Germany | 0.347277 |
| Gaza | 0.347453 |
| Seychelles | 0.350695 |
| Czechia | 0.352303 |
| Northern Mariana Is. (USA) | 0.354963 |
| Rwanda | 0.355438 |
| Clipperton Island | 0.356225 |
| Saint Lucia | 0.357872 |

| | |
|---|---|
| Sint Maarten | 0.360123 |
| Thailand | 0.364089 |
| Tokelau | 0.366644 |
| Poland | 0.367657 |
| Micronesia (Federated States of) | 0.370052 |
| Sao Tome and Principe | 0.376642 |
| Republic of Korea | 0.381915 |
| Cayman Islands | 0.38372 |
| Italy | 0.389822 |
| Bahamas | 0.389911 |
| Spratly Islands | 0.390269 |
| Ukraine | 0.396094 |
| Senkaku Islands | 0.397885 |
| Saint Martin | 0.39966 |
| Andorra | 0.405214 |
| Serbia | 0.412636 |
| Jamaica | 0.413419 |
| Hong Kong | 0.418508 |
| Tuvalu | 0.420349 |
| Wallis and Futuna | 0.420379 |
| France | 0.423442 |
| Romania | 0.424695 |
| Togo | 0.428133 |
| Slovakia | 0.428315 |
| Lebanon | 0.430933 |
| Saint Helena | 0.438642 |
| Maldives | 0.439116 |
| Uganda | 0.440196 |
| Lithuania | 0.441707 |
| Viet Nam | 0.447029 |
| Bulgaria | 0.448577 |
| Cyprus | 0.450607 |
| Belarus | 0.455567 |
| Japan | 0.459319 |
| Austria | 0.460324 |
| Marshall Islands | 0.4605 |
| United Kingdom of Great Britain & Northern Ireland | 0.463543 |
| Fiji | 0.464288 |
| Gambia | 0.466366 |
| Guinea-Bissau | 0.470239 |
| Kiribati | 0.470837 |
| Türkiye | 0.471075 |
| Palau | 0.476663 |
| Croatia | 0.479043 |
| Philippines | 0.479847 |

| | |
|---|---|
| Baker Island | 0.481823 |
| Azores Islands | 0.482204 |
| Malawi | 0.485461 |
| Eswatini | 0.48848 |
| Curaçao | 0.489099 |
| Trinidad and Tobago | 0.491474 |
| Ghana | 0.491758 |
| Slovenia | 0.499695 |
| Sri Lanka | 0.503415 |
| Barbados | 0.503876 |
| Aruba | 0.510927 |
| Benin | 0.514917 |
| Cuba | 0.518024 |
| Côte d'Ivoire | 0.518233 |
| Spain | 0.519734 |
| Malta | 0.520644 |
| Switzerland | 0.521128 |
| Åland Islands | 0.526221 |
| Taiwan | 0.527876 |
| Greece | 0.530152 |
| Bosnia and Herzegovina | 0.531287 |
| Israel | 0.532343 |
| Ireland | 0.537144 |
| Dominican Republic | 0.540215 |
| North Macedonia | 0.540325 |
| Isle of Man | 0.544213 |
| Burkina Faso | 0.545084 |
| Azerbaijan | 0.547952 |
| Albania | 0.554814 |
| Panama | 0.569449 |
| Pakistan | 0.569912 |
| Senegal | 0.574444 |
| Timor-Leste | 0.576092 |
| Costa Rica | 0.576706 |
| Cabo Verde | 0.581526 |
| Lesotho | 0.58647 |
| Nepal | 0.587846 |
| Democratic People's Republic of Korea | 0.589115 |
| Armenia | 0.59027 |
| Guatemala | 0.591362 |
| Georgia | 0.595615 |
| Cambodia | 0.597994 |
| Syrian Arab Republic | 0.599248 |
| Latvia | 0.611386 |
| Nicaragua | 0.614039 |
| Kuwait | 0.616822 |

| | |
|---|---|
| Myanmar | 0.625552 |
| Eithiopia | 0.640795 |
| Estonia | 0.641729 |
| Bahrain | 0.6488 |
| Montenegro | 0.658486 |
| China | 0.660995 |
| Indonesia | 0.663846 |
| Cocos (Keeling) Islands | 0.670823 |
| Mexico | 0.67428 |
| Guinea | 0.677442 |
| Tunisia | 0.680434 |
| Honduras | 0.681236 |
| Qatar | 0.685705 |
| Morocco | 0.687662 |
| Malaysia | 0.69105 |
| Argentina | 0.691207 |
| United Republic of Tanzania | 0.692165 |
| Kenya | 0.69338 |
| Uzbekistan | 0.694688 |
| Uruguay | 0.69585 |
| United Arab Emirates | 0.697498 |
| Sierra Leone | 0.697894 |
| Ecuador | 0.701795 |
| United States of America | 0.70627 |
| Iran (Islamic Republic of) | 0.706536 |
| Paraguay | 0.707325 |
| Paracel Islands | 0.707379 |
| Zimbabwe | 0.713448 |
| Dekelia | 0.716206 |
| French Southern Territories | 0.728943 |
| Madagascar | 0.730048 |
| Iraq | 0.731387 |
| Monaco | 0.731719 |
| Cameroon | 0.745164 |
| Finland | 0.74569 |
| Norway | 0.749206 |
| New Zealand | 0.757322 |
| Belize | 0.762496 |
| Afghanistan | 0.764118 |
| Eritrea | 0.764328 |
| South Sudan | 0.766917 |
| Somalia | 0.769584 |
| Sweden | 0.771101 |
| Lao People's Democratic Republic | 0.771156 |
| Sudan | 0.772566 |
| Kazakhstan | 0.77468 |

| | |
|---|---|
| Turkmenistan | 0.780521 |
| Jammu and Kashmir | 0.780713 |
| Kuril Islands | 0.785549 |
| Kyrgyzstan | 0.786037 |
| Tajikistan | 0.786494 |
| Zambia | 0.787105 |
| Colombia | 0.790324 |
| Chile | 0.790611 |
| Congo | 0.7918 |
| Brunei Darussalam | 0.797517 |
| Brazil | 0.802152 |
| Mozambique | 0.80526 |
| Papua New Guinea | 0.805605 |
| Abyei | 0.806413 |
| Angola | 0.808601 |
| Liberia | 0.809232 |
| Democratic Republic of the Congo | 0.809574 |
| Bolivia (Plurinational State of) | 0.810284 |
| Venezuela | 0.814841 |
| Djibouti | 0.818986 |
| Mali | 0.819409 |
| Bassas da India | 0.821552 |
| Arunachal Pradesh | 0.82314 |
| Bhutan | 0.82414 |
| Peru | 0.831756 |
| Yemen | 0.832031 |
| Equatorial Guinea | 0.837453 |
| Jordan | 0.840376 |
| Chad | 0.841708 |
| Niger | 0.844414 |
| Central African Republic | 0.852046 |
| Bouvet Island | 0.857888 |
| Gabon | 0.868986 |
| Saudi Arabia | 0.869558 |
| Iceland | 0.869815 |
| Mongolia | 0.87874 |
| Oman | 0.878951 |
| Namibia | 0.880732 |
| Russian Federation | 0.881498 |
| Australia | 0.884048 |
| Botswana | 0.89415 |
| China/India | 0.898139 |
| Canada | 0.903494 |
| Algeria | 0.918261 |
| South Georgia and the South Sandwich Islands | 0.931876 |
| Heard Island and McDonald Islands | 0.936341 |

| | |
|---|---|
| Mauritania | 0.936945 |
| Egypt | 0.937033 |
| Aksai Chin | 0.947629 |
| Guyana | 0.949829 |
| Libya | 0.951685 |
| Suriname | 0.956078 |
| French Guiana | 0.95627 |
| Western Sahara | 0.976052 |
| Svalbard and Jan Mayen Islands | 0.98357 |
| Greenland | 0.993105 |

**Table S3** shows the mean EII values per A) ecoregion and B) ecosystem.

**A)**

| Ecoregion_name | MEAN EII |
|---|---|
| Red River freshwater swamp forests | 0.11941 |
| Lower Gangetic Plains moist deciduous forests | 0.124241 |
| Upper Gangetic Plains moist deciduous forests | 0.12723 |
| Chao Phraya freshwater swamp forests | 0.128351 |
| Sundarbans freshwater swamp forests | 0.128769 |
| Windward Islands moist forests | 0.133905 |
| Po Basin mixed forests | 0.151199 |
| Mandara Plateau woodlands | 0.152074 |
| Huang He Plain mixed forests | 0.162394 |
| Veracruz dry forests | 0.171231 |
| New Caledonia rain forests | 0.172792 |
| Tubuai tropical moist forests | 0.173754 |
| Lord Howe Island subtropical forests | 0.180519 |
| Cocos Island moist forests | 0.182311 |
| Orissa semi-evergreen forests | 0.188829 |
| East Deccan dry-evergreen forests | 0.192958 |
| New Caledonia dry forests | 0.199447 |
| Vanuatu rain forests | 0.202601 |
| Cross-Niger transition forests | 0.207351 |
| Faroe Islands boreal grasslands | 0.207922 |
| Tongan tropical moist forests | 0.212454 |
| Tonle Sap-Mekong peat swamp forests | 0.214221 |
| Southeast US mixed woodlands and savannas | 0.215111 |
| Irrawaddy freshwater swamp forests | 0.217308 |
| Norfolk Island subtropical forests | 0.221222 |
| Guayaquil flooded grasslands | 0.221692 |
| Eastern Java-Bali rain forests | 0.22376 |
| Khathiar-Gir dry deciduous forests | 0.227174 |

| | |
|---|---|
| Central Deccan Plateau dry deciduous forests | 0.22798 |
| Sichuan Basin evergreen broadleaf forests | 0.229317 |
| Cook Islands tropical moist forests | 0.2321 |
| South Deccan Plateau dry deciduous forests | 0.233574 |
| Deccan thorn scrub forests | 0.236285 |
| Trinidad and Tobago dry forest | 0.239503 |
| Mascarene forests | 0.240717 |
| Comoros forests | 0.243937 |
| Society Islands tropical moist forests | 0.24545 |
| Jos Plateau forest-grassland | 0.247415 |
| Chhota-Nagpur dry deciduous forests | 0.248626 |
| Northeast China Plain deciduous forests | 0.250188 |
| Bermuda subtropical conifer forests | 0.250621 |
| Western Java rain forests | 0.251252 |
| Sierra de los Tuxtlas | 0.254015 |
| Fernando de Noronha-Atol das Rocas moist forests | 0.255594 |
| Aravalli west thorn scrub forests | 0.26034 |
| Irrawaddy dry forests | 0.26475 |
| Narmada Valley dry deciduous forests | 0.268074 |
| Aldabra Island xeric scrub | 0.273919 |
| Bahamian pineyards | 0.280344 |
| Fiji tropical dry forests | 0.287441 |
| Lesser Antillean dry forests | 0.289799 |
| Chao Phraya lowland moist deciduous forests | 0.290101 |
| English Lowlands beech forests | 0.291978 |
| Northern Vietnam lowland rain forests | 0.293377 |
| Tristan Da Cunha-Gough Islands shrub and grasslands | 0.300086 |
| Brahmaputra Valley semi-evergreen forests | 0.300245 |
| Yellow Sea saline meadow | 0.301785 |
| Malabar Coast moist forests | 0.305419 |
| Terai-Duar savanna and grasslands | 0.306222 |
| Veracruz moist forests | 0.307774 |
| St. Helena scrub and woodlands | 0.311055 |
| Humid Pampas | 0.315402 |
| Eastern Mediterranean conifer-broadleaf forests | 0.315762 |
| Northwest Hawai'i scrub | 0.316158 |
| European Atlantic mixed forests | 0.317526 |
| Jamaican dry forests | 0.318089 |
| Changjiang Plain evergreen forests | 0.318427 |
| Nenjiang River grassland | 0.321208 |
| Tonle Sap freshwater swamp forests | 0.321682 |
| Leeward Islands moist forests | 0.323176 |
| Indus Valley desert | 0.3237 |
| Samoan tropical moist forests | 0.32501 |

| | |
|---|---|
| Chiapas Depression dry forests | 0.325503 |
| Pantanos de Centla | 0.326102 |
| Puerto Rican moist forests | 0.327466 |
| Southern Great Lakes forests | 0.32945 |
| Cayos Miskitos-San Andrés and Providencia moist forests | 0.329684 |
| Greater Negros-Panay rain forests | 0.336495 |
| Indochina mangroves | 0.337822 |
| Marquesas tropical moist forests | 0.338128 |
| Puerto Rican dry forests | 0.340428 |
| North Deccan dry deciduous forests | 0.343678 |
| Central Anatolian steppe and woodlands | 0.343949 |
| Pannonian mixed forests | 0.344899 |
| Marianas tropical dry forests | 0.349856 |
| Central Indochina dry forests | 0.349891 |
| Italian sclerophyllous and semi-deciduous forests | 0.350703 |
| Caribbean shrublands | 0.35125 |
| Bohai Sea saline meadow | 0.354782 |
| Nigerian lowland forests | 0.357556 |
| Central US forest-grasslands transition | 0.358212 |
| East Deccan moist deciduous forests | 0.35836 |
| Western Java montane rain forests | 0.360005 |
| Carolines tropical moist forests | 0.360064 |
| Nile Delta flooded savanna | 0.362262 |
| Ethiopian montane moorlands | 0.362361 |
| Bajío dry forests | 0.363804 |
| Central Anatolian steppe | 0.364448 |
| South Western Ghats moist deciduous forests | 0.364582 |
| North Western Ghats moist deciduous forests | 0.369713 |
| Baltic mixed forests | 0.370387 |
| Ethiopian montane grasslands and woodlands | 0.371428 |
| Panamanian dry forests | 0.372016 |
| Godavari-Krishna mangroves | 0.372833 |
| Central Polynesian tropical moist forests | 0.375299 |
| Eastern Guinean forests | 0.376252 |
| Central Korean deciduous forests | 0.377184 |
| Taiheiyo evergreen forests | 0.379261 |
| Victoria Basin forest-savanna | 0.381878 |
| Kermadec Islands subtropical moist forests | 0.383118 |
| Christmas and Cocos Islands tropical forests | 0.383245 |
| Northern Khorat Plateau moist deciduous forests | 0.383723 |
| Central Tallgrass prairie | 0.384275 |
| Eastern Java-Bali montane rain forests | 0.390337 |
| California Central Valley grasslands | 0.392193 |
| Hispaniolan moist forests | 0.39432 |

| | |
|---|---|
| Southern Korea evergreen forests | 0.395438 |
| Northern Tallgrass prairie | 0.396691 |
| Southern Vietnam lowland dry forests | 0.401391 |
| Western European broadleaf forests | 0.401439 |
| Central European mixed forests | 0.401811 |
| Tuamotu tropical moist forests | 0.402071 |
| East European forest steppe | 0.40275 |
| Sumatran freshwater swamp forests | 0.403169 |
| São Tomé, Príncipe, and Annobón forests | 0.404539 |
| South China-Vietnam subtropical evergreen forests | 0.405772 |
| Niger Delta swamp forests | 0.410307 |
| Balkan mixed forests | 0.41184 |
| Himalayan subtropical pine forests | 0.412078 |
| Maldives-Lakshadweep-Chagos Archipelago tropical moist forests | 0.412193 |
| South Western Ghats montane rain forests | 0.413136 |
| East African montane forests | 0.414209 |
| Nihonkai evergreen forests | 0.415 |
| Sundarbans mangroves | 0.416981 |
| Northeast US Coastal forests | 0.418754 |
| Tyrrhenian-Adriatic sclerophyllous and mixed forests | 0.419976 |
| Mississippi lowland forests | 0.420767 |
| Ile Europa and Bassas da India xeric scrub | 0.421843 |
| Cauca Valley dry forests | 0.423098 |
| Pontic steppe | 0.430781 |
| Mediterranean woodlands and forests | 0.432026 |
| Tehuacán Valley matorral | 0.433499 |
| Canadian Aspen forests and parklands | 0.43367 |
| Kwazulu Natal-Cape coastal forests | 0.434493 |
| Jamaican moist forests | 0.435827 |
| Myanmar Coast mangroves | 0.436139 |
| Yap tropical dry forests | 0.438545 |
| Luzon rain forests | 0.441617 |
| Atlantic coastal pine barrens | 0.442682 |
| Mount Cameroon and Bioko montane forests | 0.45161 |
| Willamette Valley oak savanna | 0.452721 |
| Central China Loess Plateau mixed forests | 0.454408 |
| West Sudanian savanna | 0.455229 |
| Anatolian conifer and deciduous mixed forests | 0.455783 |
| Cyprus Mediterranean forests | 0.457091 |
| Hispaniolan dry forests | 0.457278 |
| Eastern Great Lakes lowland forests | 0.457809 |
| Granitic Seychelles forests | 0.458889 |
| Irrawaddy moist deciduous forests | 0.459839 |
| Southeast Iberian shrubs and woodlands | 0.459953 |

| | |
|---|---|
| Cuban dry forests | 0.460178 |
| Sri Lanka lowland rain forests | 0.460275 |
| Eastern Micronesia tropical moist forests | 0.461844 |
| Celtic broadleaf forests | 0.463013 |
| Mindanao-Eastern Visayas rain forests | 0.465476 |
| Upper Midwest US forest-savanna transition | 0.465554 |
| Alto Paraná Atlantic forests | 0.466387 |
| Myanmar coastal rain forests | 0.468456 |
| Daba Mountains evergreen forests | 0.470437 |
| Southwest Iberian Mediterranean sclerophyllous and mixed forests | 0.471447 |
| Veracruz montane forests | 0.471614 |
| Taiheiyo montane deciduous forests | 0.473573 |
| Isthmian-Pacific moist forests | 0.474374 |
| Aegean and Western Turkey sclerophyllous and mixed forests | 0.4767 |
| Caspian Hyrcanian mixed forests | 0.476767 |
| Central American dry forests | 0.478004 |
| Himalayan subtropical broadleaf forests | 0.479306 |
| Central-Southern US mixed grasslands | 0.479871 |
| Western Gulf coastal grasslands | 0.482993 |
| Texas blackland prairies | 0.483523 |
| Iberian sclerophyllous and semi-deciduous forests | 0.483646 |
| Palau tropical moist forests | 0.483918 |
| Azerbaijan shrub desert and steppe | 0.486044 |
| Mediterranean conifer and mixed forests | 0.486608 |
| Southern Pacific dry forests | 0.487018 |
| Juan Fernández Islands temperate forests | 0.487778 |
| Tenasserim-South Thailand semi-evergreen rain forests | 0.488268 |
| North Western Ghats montane rain forests | 0.489411 |
| Peninsular Malaysian rain forests | 0.49024 |
| Western Himalayan subalpine conifer forests | 0.492261 |
| Motagua Valley thornscrub | 0.494517 |
| Guizhou Plateau broadleaf and mixed forests | 0.49487 |
| Central Mexican matorral | 0.495649 |
| Nihonkai montane deciduous forests | 0.495826 |
| Eastern Anatolian deciduous forests | 0.496447 |
| Fiji tropical moist forests | 0.496743 |
| Costa Rican seasonal moist forests | 0.498487 |
| Trinidad and Tobago moist forest | 0.499566 |
| Sri Lanka montane rain forests | 0.500262 |
| Western Himalayan broadleaf forests | 0.504226 |
| Eastern Anatolian montane steppe | 0.505464 |
| Sumatran lowland rain forests | 0.506927 |
| South Apennine mixed montane forests | 0.50772 |
| Eyre and York mallee | 0.509063 |

| | |
|---|---|
| Suiphun-Khanka meadows and forest meadows | 0.510357 |
| Mindoro rain forests | 0.51225 |
| Sumatran peat swamp forests | 0.51303 |
| Euxine-Colchic broadleaf forests | 0.513104 |
| Sri Lanka dry-zone dry evergreen forests | 0.514611 |
| Cantabrian mixed forests | 0.517372 |
| Sierra Madre de Chiapas moist forests | 0.518896 |
| California coastal sage and chaparral | 0.51897 |
| Hawai'i tropical low shrublands | 0.520414 |
| Northeast Spain and Southern France Mediterranean forests | 0.520737 |
| Jian Nan subtropical evergreen forests | 0.520791 |
| Thar desert | 0.521173 |
| Northern Mesoamerican Pacific mangroves | 0.521749 |
| Central African mangroves | 0.521809 |
| Sulawesi lowland rain forests | 0.522263 |
| Illyrian deciduous forests | 0.522602 |
| Scandinavian coastal conifer forests | 0.523246 |
| Alai-Western Tian Shan steppe | 0.524388 |
| South Taiwan monsoon rain forests | 0.524461 |
| Petén-Veracruz moist forests | 0.524845 |
| Meghalaya subtropical forests | 0.525665 |
| Sulu Archipelago rain forests | 0.528422 |
| South China Sea Islands | 0.528718 |
| Aleutian Islands tundra | 0.529343 |
| Taiwan subtropical evergreen forests | 0.529465 |
| Yunnan Plateau subtropical evergreen forests | 0.530999 |
| Ogasawara subtropical moist forests | 0.531248 |
| Zagros Mountains forest steppe | 0.531821 |
| Bahamian-Antillean mangroves | 0.532516 |
| Guinean forest-savanna | 0.532663 |
| Jalisco dry forests | 0.534183 |
| Magdalena-Urabá moist forests | 0.534695 |
| Appalachian Piedmont forests | 0.536456 |
| Maracaibo dry forests | 0.541356 |
| Southern Africa mangroves | 0.54158 |
| Western Polynesian tropical moist forests | 0.542108 |
| Hainan Island monsoon rain forests | 0.543816 |
| Guinean mangroves | 0.544455 |
| Ascension scrub and grasslands | 0.545463 |
| Knysna-Amatole montane forests | 0.547573 |
| Isthmian-Atlantic moist forests | 0.547677 |
| Trans-Mexican Volcanic Belt pine-oak forests | 0.548048 |
| Renosterveld shrubland | 0.548455 |
| Mid-Atlantic US coastal savannas | 0.550004 |

| | |
|---|---|
| Cuban cactus scrub | 0.552562 |
| Appenine deciduous montane forests | 0.552823 |
| Tamaulipan matorral | 0.553314 |
| Balsas dry forests | 0.555414 |
| Southeast Indochina dry evergreen forests | 0.558001 |
| Ordos Plateau steppe | 0.558236 |
| Canary Islands dry woodlands and forests | 0.558822 |
| Honshu alpine conifer forests | 0.559102 |
| Albertine Rift montane forests | 0.559437 |
| Northern Anatolian conifer and deciduous forests | 0.559488 |
| Cape Verde Islands dry forests | 0.560445 |
| Sumba deciduous forests | 0.561996 |
| Lesser Sundas deciduous forests | 0.562751 |
| Ethiopian montane forests | 0.562809 |
| Chiapas montane forests | 0.564671 |
| Hokkaido deciduous forests | 0.565188 |
| Peninsular Malaysian peat swamp forests | 0.566647 |
| Interior Plateau US Hardwood Forests | 0.567135 |
| Ecuadorian dry forests | 0.567483 |
| Western Ecuador moist forests | 0.567514 |
| Espinal | 0.568817 |
| Pernambuco interior forests | 0.568938 |
| Indus River Delta-Arabian Sea mangroves | 0.569828 |
| Southern Mesoamerican Pacific mangroves | 0.571551 |
| Pernambuco coastal forests | 0.571965 |
| Enriquillo wetlands | 0.575769 |
| Crete Mediterranean forests | 0.575872 |
| Cauca Valley montane forests | 0.576121 |
| Everglades flooded grasslands | 0.576789 |
| Highveld grasslands | 0.578049 |
| Guinean montane forests | 0.578766 |
| Amur meadow steppe | 0.579957 |
| Qin Ling Mountains deciduous forests | 0.582273 |
| Central American pine-oak forests | 0.58329 |
| Atlantic Coast restingas | 0.583399 |
| Timor and Wetar deciduous forests | 0.584257 |
| Northwest Iberian montane forests | 0.584859 |
| Yucatán dry forests | 0.586102 |
| Alps conifer and mixed forests | 0.586245 |
| Palawan rain forests | 0.586323 |
| Cameroon Highlands forests | 0.586852 |
| Southwest Arabian montane woodlands and grasslands | 0.587096 |
| Oaxacan montane forests | 0.587904 |
| Drakensberg Escarpment savanna and thicket | 0.587977 |

| | |
|---|---|
| Paraná flooded savanna | 0.589052 |
| Southern Acacia-Commiphora bushlands and thickets | 0.590996 |
| Carpathian montane forests | 0.59244 |
| Cuban moist forests | 0.592788 |
| Sinú Valley dry forests | 0.593923 |
| Mesoamerican Gulf-Caribbean mangroves | 0.593942 |
| Caatinga Enclaves moist forests | 0.594278 |
| Rwenzori-Virunga montane moorlands | 0.597062 |
| Guajira-Barranquilla xeric scrub | 0.598073 |
| Syrian xeric grasslands and shrublands | 0.598766 |
| Hokkaido montane conifer forests | 0.601453 |
| Southern Anatolian montane conifer and deciduous forests | 0.601905 |
| Madagascar ericoid thickets | 0.602625 |
| Southwest Australia savanna | 0.603807 |
| Hispaniolan pine forests | 0.605244 |
| Magdalena Valley dry forests | 0.605372 |
| Palouse prairie | 0.608464 |
| Hawai'i tropical dry forests | 0.610345 |
| Socotra Island xeric shrublands | 0.610365 |
| Central American montane forests | 0.613075 |
| Tigris-Euphrates alluvial salt marsh | 0.615397 |
| Southeast US conifer savannas | 0.615818 |
| Mulanje Montane forest-grassland | 0.618328 |
| Kazakh steppe | 0.618693 |
| Andaman Islands rain forests | 0.619345 |
| Southern Rift Montane forest-grassland | 0.619549 |
| Madeira evergreen forests | 0.621186 |
| Southern Atlantic Brazilian mangroves | 0.622928 |
| Azores temperate mixed forests | 0.623552 |
| Appalachian-Blue Ridge forests | 0.623621 |
| Sarmatic mixed forests | 0.625338 |
| Catatumbo moist forests | 0.625741 |
| Cuban pine forests | 0.627952 |
| Central Andean wet puna | 0.628328 |
| Halmahera rain forests | 0.628376 |
| East African mangroves | 0.628666 |
| Northern Shortgrass prairie | 0.630081 |
| Elburz Range forest steppe | 0.631714 |
| Pindus Mountains mixed forests | 0.631919 |
| Serra do Mar coastal forests | 0.632258 |
| East Central Texas forests | 0.632502 |
| Crimean Submediterranean forest complex | 0.633234 |
| Kazakh forest steppe | 0.636628 |
| Cordillera La Costa montane forests | 0.636722 |

| | |
|---|---|
| Northern Swahili coastal forests | 0.637907 |
| Western shortgrass prairie | 0.640176 |
| La Costa xeric shrublands | 0.640699 |
| Chatham Island temperate forests | 0.641528 |
| Rodope montane mixed forests | 0.643225 |
| Iberian conifer forests | 0.643511 |
| Lake Chad flooded savanna | 0.643863 |
| Southeast Australia temperate forests | 0.643884 |
| Sulawesi montane rain forests | 0.644158 |
| Northern Indochina subtropical forests | 0.644299 |
| Magdalena Valley montane forests | 0.644935 |
| Marañón dry forests | 0.64543 |
| Humid Chaco | 0.648235 |
| Sunda Shelf mangroves | 0.64849 |
| Mediterranean Acacia-Argania dry woodlands and succulent thickets | 0.64939 |
| Puget lowland forests | 0.650271 |
| Sumatran montane rain forests | 0.651606 |
| Dinaric Mountains mixed forests | 0.651659 |
| Sinaloan dry forests | 0.651705 |
| Caucasus mixed forests | 0.653478 |
| Serengeti volcanic grasslands | 0.654799 |
| Paropamisus xeric woodlands | 0.654945 |
| Gissaro-Alai open woodlands | 0.655433 |
| South American Pacific mangroves | 0.655537 |
| Northwest Andean montane forests | 0.655944 |
| Cross-Timbers savanna-woodland | 0.6566 |
| North Atlantic moist mixed forests | 0.656806 |
| Mindanao montane rain forests | 0.65682 |
| Maputaland coastal forests and woodlands | 0.657218 |
| California interior chaparral and woodlands | 0.657717 |
| Venezuelan Andes montane forests | 0.658051 |
| Southwest Arabian coastal xeric shrublands | 0.65844 |
| Helanshan montane conifer forests | 0.658771 |
| Cardamom Mountains rain forests | 0.659306 |
| Araya and Paria xeric scrub | 0.659333 |
| Limpopo lowveld | 0.661546 |
| Drakensberg grasslands | 0.663083 |
| Fynbos shrubland | 0.664432 |
| Southwest Arabian Escarpment shrublands and woodlands | 0.664618 |
| Sonoran-Sinaloan subtropical dry forest | 0.664682 |
| Piney Woods | 0.666586 |
| Biak-Numfoor rain forests | 0.66664 |
| Banda Sea Islands moist deciduous forests | 0.668175 |
| Southwest Australia woodlands | 0.670451 |

| | |
|---|---|
| Antipodes Subantarctic Islands tundra | 0.670912 |
| Flint Hills tallgrass prairie | 0.670987 |
| Central American Atlantic moist forests | 0.671288 |
| Northern Thailand-Laos moist deciduous forests | 0.67144 |
| Seram rain forests | 0.671455 |
| Araucaria moist forests | 0.672781 |
| Esperance mallee | 0.672802 |
| Angolan montane forest-grassland | 0.673221 |
| Kopet Dag semi-desert | 0.673702 |
| Bolivian montane dry forests | 0.67414 |
| Paraguaná xeric scrub | 0.674444 |
| Uruguayan savanna | 0.674536 |
| Hawai'i tropical moist forests | 0.675311 |
| Central Asian riparian woodlands | 0.677022 |
| San Lucan xeric scrub | 0.677139 |
| Southwest Arabian highland xeric scrub | 0.677519 |
| Louisiade Archipelago rain forests | 0.678236 |
| Sundaland heath forests | 0.678284 |
| Manchurian mixed forests | 0.67975 |
| Ozark Mountain forests | 0.680224 |
| Badghyz and Karabil semi-desert | 0.682577 |
| Pyrenees conifer and mixed forests | 0.682756 |
| Magellanic subpolar forests | 0.682894 |
| Madagascar subhumid forests | 0.683872 |
| Appalachian mixed mesophytic forests | 0.684119 |
| Apure-Villavicencio dry forests | 0.685868 |
| Nyanga-Chimanimani Montane forest-grassland | 0.686789 |
| Buru rain forests | 0.688453 |
| Tamaulipan mezquital | 0.690817 |
| Central bushveld | 0.69089 |
| Nansei Islands subtropical evergreen forests | 0.691413 |
| Lara-Falcón dry forests | 0.691775 |
| Eastern Arc forests | 0.69206 |
| Nicobar Islands rain forests | 0.69237 |
| Northern Andean páramo | 0.692496 |
| Cordillera Central páramo | 0.693778 |
| Northland temperate kauri forests | 0.693839 |
| Bahia coastal forests | 0.694189 |
| Angolan scarp savanna and woodlands | 0.695132 |
| Ozark Highlands mixed forests | 0.695138 |
| Southwest Borneo freshwater swamp forests | 0.696526 |
| Southeast Australia temperate savanna | 0.696582 |
| Southern Cone Mesopotamian savanna | 0.696802 |
| Western Guinean lowland forests | 0.697653 |

| | |
|---|---|
| Baluchistan xeric woodlands | 0.697702 |
| Allegheny Highlands forests | 0.697944 |
| Mongolian-Manchurian grassland | 0.698325 |
| Zambezian-Limpopo mixed woodlands | 0.699017 |
| Eastern Panamanian montane forests | 0.699748 |
| Cross-Sanaga-Bioko coastal forests | 0.700044 |
| Nebraska Sand Hills mixed grasslands | 0.702882 |
| Qionglai-Minshan conifer forests | 0.7052 |
| Chilean Matorral | 0.705543 |
| Tumbes-Piura dry forests | 0.706443 |
| Amazon-Orinoco-Southern Caribbean mangroves | 0.707231 |
| Itigi-Sumbu thicket | 0.70835 |
| Santa Marta páramo | 0.708555 |
| East Sudanian savanna | 0.708765 |
| Canterbury-Otago tussock grasslands | 0.709192 |
| Bahia interior forests | 0.71005 |
| Solomon Islands rain forests | 0.710246 |
| Luzon tropical pine forests | 0.710318 |
| South Siberian forest steppe | 0.711319 |
| Mizoram-Manipur-Kachin rain forests | 0.711637 |
| Inner Niger Delta flooded savanna | 0.712075 |
| Dry Chaco | 0.71259 |
| Peruvian Yungas | 0.715607 |
| Northeast India-Myanmar pine forests | 0.716589 |
| East African montane moorlands | 0.717014 |
| Naracoorte woodlands | 0.717362 |
| Al-Hajar foothill xeric woodlands and shrublands | 0.717933 |
| Montana Valley and Foothill grasslands | 0.718618 |
| Sumatran tropical pine forests | 0.718726 |
| Sulaiman Range alpine meadows | 0.71877 |
| Southern Annamites montane rain forests | 0.718978 |
| Eastern Cordillera Real montane forests | 0.719737 |
| Trobriand Islands rain forests | 0.720431 |
| Cordillera de Merida páramo | 0.722286 |
| Qilian Mountains conifer forests | 0.722799 |
| Chimalapas montane forests | 0.72369 |
| Northern Acacia-Commiphora bushlands and thickets | 0.724269 |
| Changbai Mountains mixed forests | 0.72479 |
| Borneo peat swamp forests | 0.725293 |
| Talamancan montane forests | 0.725405 |
| Al-Hajar montane woodlands and shrublands | 0.726268 |
| Etosha Pan halophytics | 0.727608 |
| Tian Shan foothill arid steppe | 0.728184 |
| Cerrado | 0.728402 |

| | |
|---|---|
| Sierra Madre del Sur pine-oak forests | 0.729018 |
| Kopet Dag woodlands and forest steppe | 0.729321 |
| Cuban wetlands | 0.729996 |
| Mentawai Islands rain forests | 0.730202 |
| Luang Prabang montane rain forests | 0.730852 |
| East Afghan montane conifer forests | 0.730942 |
| Madagascar spiny thickets | 0.731076 |
| Santa Marta montane forests | 0.732323 |
| Albany thickets | 0.734869 |
| Kazakh upland steppe | 0.734948 |
| Eastern Himalayan broadleaf forests | 0.735239 |
| Sierra Madre de Oaxaca pine-oak forests | 0.73544 |
| Yapen rain forests | 0.735627 |
| Chiquitano dry forests | 0.736925 |
| Southern Andean Yungas | 0.737335 |
| Sierra de la Laguna dry forests | 0.737479 |
| Caledon conifer forests | 0.738762 |
| Rapa Nui and Sala y Gómez subtropical forests | 0.738984 |
| Caatinga | 0.740772 |
| Emin Valley steppe | 0.743295 |
| Campos Rupestres montane savanna | 0.744137 |
| Corsican montane broadleaf and mixed forests | 0.744793 |
| Murray-Darling woodlands and mallee | 0.745265 |
| Patía valley dry forests | 0.746182 |
| Gurupa várzea | 0.746428 |
| Kayah-Karen montane rain forests | 0.747206 |
| Madagascar dry deciduous forests | 0.748967 |
| Cordillera Oriental montane forests | 0.751568 |
| East African halophytics | 0.753668 |
| Chocó-Darién moist forests | 0.754847 |
| Valdivian temperate forests | 0.754979 |
| Tasmanian temperate forests | 0.755056 |
| Alaska Peninsula montane taiga | 0.756436 |
| Central Zambezian wet miombo woodlands | 0.758531 |
| Madagascar humid forests | 0.75877 |
| Tian Shan montane conifer forests | 0.761303 |
| Northern Pacific Alaskan coastal forests | 0.761881 |
| Hengduan Mountains subalpine conifer forests | 0.762053 |
| Northern Annamites rain forests | 0.763748 |
| Altai steppe and semi-desert | 0.763755 |
| Madagascar mangroves | 0.763963 |
| Meseta Central matorral | 0.764111 |
| New Zealand South Island montane grasslands | 0.764263 |
| Central Afghan Mountains xeric woodlands | 0.76478 |

| | |
|---|---|
| Mediterranean dry woodlands and steppe | 0.765299 |
| New Zealand North Island temperate forests | 0.766355 |
| Somali Acacia-Commiphora bushlands and thickets | 0.767173 |
| Trans Fly savanna and grasslands | 0.769605 |
| Madagascar succulent woodlands | 0.76994 |
| Djibouti xeric shrublands | 0.770314 |
| Southern Swahili coastal forests and woodlands | 0.770384 |
| Northeast Brazil restingas | 0.770964 |
| Borneo lowland rain forests | 0.771418 |
| Dry miombo woodlands | 0.773414 |
| Rann of Kutch seasonal salt marsh | 0.774305 |
| Southern Congolian forest-savanna | 0.775309 |
| Arabian-Persian Gulf coastal plain desert | 0.776021 |
| New Zealand South Island temperate forests | 0.777347 |
| Western Congolian forest-savanna | 0.778199 |
| Gulf of St. Lawrence lowland forests | 0.779104 |
| Huon Peninsula montane rain forests | 0.779279 |
| South Arabian fog woodlands, shrublands, and dune | 0.779378 |
| Kinabalu montane alpine meadows | 0.77952 |
| Snake-Columbia shrub steppe | 0.779804 |
| Angolan mopane woodlands | 0.780903 |
| Zambezian coastal flooded savanna | 0.781437 |
| Belizian pine savannas | 0.782503 |
| Mediterranean High Atlas juniper steppe | 0.782686 |
| Brazilian Atlantic dry forests | 0.782858 |
| South Iran Nubo-Sindian desert and semi-desert | 0.783083 |
| Northwestern Himalayan alpine shrub and meadows | 0.783192 |
| Nujiang Langcang Gorge alpine conifer and mixed forests | 0.783443 |
| Yucatán moist forests | 0.784375 |
| Pantanal | 0.785232 |
| Sechura desert | 0.785346 |
| Papuan Central Range sub-alpine grasslands | 0.786821 |
| Maranhão Babaçu forests | 0.787159 |
| Sahelian Acacia savanna | 0.78744 |
| Southeast Papuan rain forests | 0.787944 |
| Chin Hills-Arakan Yoma montane forests | 0.788149 |
| Peninsular Malaysian montane rain forests | 0.788771 |
| Sudd flooded grasslands | 0.790419 |
| Western Congolian swamp forests | 0.790505 |
| Northern California coastal forests | 0.790888 |
| Admiralty Islands lowland rain forests | 0.791416 |
| Luzon montane rain forests | 0.791737 |
| Eastern Australian temperate forests | 0.792148 |
| Flinders-Lofty montane woodlands | 0.792593 |

| | |
|---|---|
| Western Siberian hemiboreal forests | 0.792895 |
| Yarlung Zanbo arid steppe | 0.793187 |
| Zambezian Baikiaea woodlands | 0.794339 |
| New Guinea mangroves | 0.796817 |
| Santa Lucia Montane Chaparral & Woodlands | 0.797094 |
| Tocantins/Pindare moist forests | 0.797305 |
| Central Asian southern desert | 0.798649 |
| Mesopotamian shrub desert | 0.799438 |
| Hindu Kush alpine meadow | 0.800029 |
| Afghan Mountains semi-desert | 0.800032 |
| Jarrah-Karri forest and shrublands | 0.801855 |
| Central Persian desert basins | 0.802389 |
| New Britain-New Ireland lowland rain forests | 0.803084 |
| Llanos | 0.803203 |
| Ghorat-Hazarajat alpine meadow | 0.803408 |
| Richmond temperate forests | 0.803448 |
| Sierra Madre Oriental pine-oak forests | 0.803748 |
| Edwards Plateau savanna | 0.804542 |
| Orinoco wetlands | 0.806427 |
| Hobyo grasslands and shrublands | 0.807317 |
| Southeast Tibet shrublands and meadows | 0.808759 |
| California montane chaparral and woodlands | 0.812011 |
| Caspian lowland desert | 0.812199 |
| Kuh Rud and Eastern Iran montane woodlands | 0.812857 |
| Rakiura Island temperate forests | 0.813076 |
| Northern Congolian Forest-Savanna | 0.814421 |
| Mid-Canada Boreal Plains forests | 0.814758 |
| Southern Indian Ocean Islands tundra | 0.81482 |
| Junggar Basin semi-desert | 0.815496 |
| Beni savanna | 0.815809 |
| Sierra de la Laguna pine-oak forests | 0.816012 |
| Sierra Madre Occidental pine-oak forests | 0.81721 |
| North Arabian highland shrublands | 0.817221 |
| Guianan freshwater swamp forests | 0.817508 |
| Central Range Papuan montane rain forests | 0.817583 |
| Marajó várzea | 0.817907 |
| Tian Shan montane steppe and meadows | 0.817932 |
| Beringia lowland tundra | 0.81904 |
| Horn of Africa xeric bushlands | 0.820889 |
| Tasmanian Central Highland forests | 0.821401 |
| Zambezian mopane woodlands | 0.821555 |
| Northeast Congolian lowland forests | 0.821812 |
| New England-Acadian forests | 0.821928 |
| Northern New Guinea lowland rain and freshwater swamp forests | 0.822137 |

| | |
|---|---|
| Angolan wet miombo woodlands | 0.822865 |
| Central Andean puna | 0.823424 |
| Eastern Himalayan subalpine conifer forests | 0.825648 |
| Scandinavian Montane Birch forest and grasslands | 0.827365 |
| Southern New Guinea freshwater swamp forests | 0.828319 |
| Brigalow tropical savanna | 0.833098 |
| Queensland tropical rain forests | 0.835368 |
| Western Great Lakes forests | 0.837144 |
| Mato Grosso tropical dry forests | 0.837705 |
| Vogelkop-Aru lowland rain forests | 0.837712 |
| Red Sea-Arabian Desert shrublands | 0.838246 |
| Red Sea mangroves | 0.839516 |
| Wyoming Basin shrub steppe | 0.839802 |
| Congolian coastal forests | 0.84067 |
| Blue Mountains forests | 0.840679 |
| Sonoran desert | 0.840941 |
| Eastern Cascades forests | 0.84187 |
| Zambezian flooded grasslands | 0.842857 |
| Monte Alegre várzea | 0.842906 |
| Wasatch and Uinta montane forests | 0.844473 |
| Gulf of California xeric scrub | 0.845823 |
| Selenge-Orkhon forest steppe | 0.846139 |
| Succulent Karoo xeric shrublands | 0.846368 |
| Qilian Mountains subalpine meadows | 0.847542 |
| Eastern Congolian swamp forests | 0.848182 |
| Daurian forest steppe | 0.848387 |
| Scandinavian and Russian taiga | 0.848523 |
| Northeast Himalayan subalpine conifer forests | 0.84931 |
| Chihuahuan desert | 0.849978 |
| Orinoco Delta swamp forests | 0.850368 |
| Central Asian northern desert | 0.850857 |
| Nama Karoo shrublands | 0.851168 |
| Tibetan Plateau alpine shrublands and meadows | 0.852604 |
| Altai montane forest and forest steppe | 0.853905 |
| Khangai Mountains alpine meadow | 0.855211 |
| Low Monte | 0.855785 |
| Iceland boreal birch forests and alpine tundra | 0.855866 |
| Xingu-Tocantins-Araguaia moist forests | 0.856255 |
| Bolivian Yungas | 0.856567 |
| Altai alpine meadow and tundra | 0.8568 |
| Patagonian steppe | 0.857078 |
| Somali montane xeric woodlands | 0.857945 |
| Kazakh semi-desert | 0.859573 |
| Northwest Congolian lowland forests | 0.860897 |

| | |
|---|---|
| Fiordland temperate forests | 0.861037 |
| British Columbia coastal conifer forests | 0.861521 |
| Arctic coastal tundra | 0.862032 |
| New Britain-New Ireland montane rain forests | 0.862373 |
| Khangai Mountains conifer forests | 0.862802 |
| Eastern Canadian Shield taiga | 0.863153 |
| Vogelkop montane rain forests | 0.863967 |
| Registan-North Pakistan sandy desert | 0.864084 |
| Miskito pine forests | 0.864558 |
| Arabian desert | 0.864972 |
| Great Lakes Basin desert steppe | 0.86537 |
| Western Himalayan alpine shrub and meadows | 0.865937 |
| Galápagos Islands xeric scrub | 0.866713 |
| Central Pacific Northwest coastal forests | 0.867502 |
| Tasmanian temperate rain forests | 0.868247 |
| Sakhalin Island taiga | 0.868495 |
| Ucayali moist forests | 0.868909 |
| North Arabian desert | 0.868958 |
| Beringia upland tundra | 0.871377 |
| Eastern Gobi desert steppe | 0.871587 |
| Namibian savanna woodlands | 0.872001 |
| Sayan Intermontane steppe | 0.872217 |
| Queen Charlotte Islands conifer forests | 0.872295 |
| Colorado Rockies forests | 0.873229 |
| Northern Triangle subtropical forests | 0.873553 |
| Urals montane forest and taiga | 0.87383 |
| Kalahari xeric savanna | 0.874369 |
| Sierra Nevada forests | 0.874604 |
| Eritrean coastal desert | 0.876101 |
| Eastern Himalayan alpine shrub and meadows | 0.876228 |
| Ahklun and Kilbuck Upland Tundra | 0.876631 |
| Klamath-Siskiyou forests | 0.876947 |
| Cook Inlet taiga | 0.877061 |
| Baja California desert | 0.877982 |
| Arctic foothills tundra | 0.878078 |
| Masai xeric grasslands and shrublands | 0.880268 |
| Tarim Basin deciduous forests and steppe | 0.881097 |
| Zambezian evergreen dry forests | 0.881295 |
| Hawai'i tropical high shrublands | 0.881433 |
| Nelson Coast temperate forests | 0.881958 |
| North Cascades conifer forests | 0.885044 |
| Gobi Lakes Valley desert steppe | 0.887505 |
| Okanogan dry forests | 0.888153 |
| Taklimakan desert | 0.888228 |

| | |
|---|---|
| Borneo montane rain forests | 0.889558 |
| Central Andean dry puna | 0.890958 |
| Kola Peninsula tundra | 0.891751 |
| Madeira-Tapajós moist forests | 0.892022 |
| Central Congolian lowland forests | 0.892639 |
| Guianan savanna | 0.893865 |
| East Arabian fog shrublands and sand desert | 0.894506 |
| Kalahari Acacia woodlands | 0.894522 |
| Central Tibetan Plateau alpine steppe | 0.895816 |
| Nullarbor Plains xeric shrublands | 0.895931 |
| Copper Plateau taiga | 0.895987 |
| Namaqualand-Richtersveld steppe | 0.896358 |
| Australian Alps montane grasslands | 0.896365 |
| Kimberly tropical savanna | 0.896939 |
| South Central Rockies forests | 0.897005 |
| High Monte | 0.898322 |
| Novosibirsk Islands Arctic desert | 0.898828 |
| Karakoram-West Tibetan Plateau alpine steppe | 0.898983 |
| Iquitos várzea | 0.900544 |
| Central-Southern Cascades Forests | 0.902009 |
| Colorado Plateau shrublands | 0.902106 |
| Westland temperate forests | 0.902284 |
| Northern New Guinea montane rain forests | 0.902348 |
| Makgadikgadi halophytics | 0.903559 |
| Alberta-British Columbia foothills forests | 0.904124 |
| Pilbara shrublands | 0.904178 |
| Southern New Guinea lowland rain forests | 0.90446 |
| Eastern Canadian forests | 0.905748 |
| Arabian sand desert | 0.905977 |
| Northern Rockies conifer forests | 0.906021 |
| Great Basin shrub steppe | 0.90603 |
| Wrangel Island Arctic desert | 0.906459 |
| Saharan halophytics | 0.906461 |
| Gibson desert | 0.907474 |
| Alashan Plateau semi-desert | 0.90804 |
| Pacific Coastal Mountain icefields and tundra | 0.90849 |
| Northern Triangle temperate forests | 0.908533 |
| Pamir alpine desert and tundra | 0.910059 |
| Victoria Plains tropical savanna | 0.910241 |
| Trans-Baikal conifer forests | 0.910343 |
| West Siberian taiga | 0.910585 |
| Brooks-British Range tundra | 0.910699 |
| South Arabian plains and plateau desert | 0.911325 |
| Sayan alpine meadows and tundra | 0.913164 |

| | |
|---|---|
| Sayan montane conifer forests | 0.914069 |
| Ussuri broadleaf and mixed forests | 0.915295 |
| Arizona Mountains forests | 0.917809 |
| Napo moist forests | 0.918355 |
| East Saharan montane xeric woodlands | 0.918747 |
| Southern Andean steppe | 0.919072 |
| Eastern Canadian Forest-Boreal transition | 0.919119 |
| Mojave desert | 0.919654 |
| Great Sandy-Tanami desert | 0.920643 |
| Scotia Sea Islands tundra | 0.921617 |
| Yamal-Gydan tundra | 0.921635 |
| Carpentaria tropical savanna | 0.922004 |
| Mitchell Grass Downs | 0.922836 |
| Atacama desert | 0.92394 |
| Tapajós-Xingu moist forests | 0.924041 |
| Northwest Russian-Novaya Zemlya tundra | 0.924752 |
| Southwest Amazon moist forests | 0.9255 |
| Gariep Karoo | 0.926781 |
| Qaidam Basin semi-desert | 0.928694 |
| Great Basin montane forests | 0.931245 |
| Kamchatka-Kurile meadows and sparse forests | 0.93151 |
| Caqueta moist forests | 0.93273 |
| Northwest Territories taiga | 0.933497 |
| Central Canadian Shield forests | 0.934189 |
| Fraser Plateau and Basin conifer forests | 0.93427 |
| Einasleigh upland savanna | 0.935771 |
| Hampton mallee and woodlands | 0.937568 |
| Midwest Canadian Shield forests | 0.938093 |
| Northern Canadian Shield taiga | 0.939017 |
| North Saharan Xeric Steppe and Woodland | 0.939226 |
| North Tibetan Plateau-Kunlun Mountains alpine desert | 0.940977 |
| Chukchi Peninsula tundra | 0.941827 |
| Tirari-Sturt stony desert | 0.942518 |
| Alaska-St. Elias Range tundra | 0.942977 |
| Arnhem Land tropical savanna | 0.944257 |
| Central British Columbia Mountain forests | 0.944515 |
| Purus várzea | 0.945141 |
| Da Hinggan-Dzhagdy Mountains conifer forests | 0.945985 |
| Northeast Siberian coastal tundra | 0.946996 |
| Coolgardie woodlands | 0.948863 |
| Interior Alaska-Yukon lowland taiga | 0.951633 |
| Central Ranges xeric scrub | 0.951739 |
| Rio Negro campinarana | 0.952276 |
| Eastern Australia mulga shrublands | 0.952484 |

| | |
|---|---|
| Purus-Madeira moist forests | 0.953094 |
| Guianan piedmont moist forests | 0.953353 |
| Uatumã-Trombetas moist forests | 0.954085 |
| Northern Cordillera forests | 0.955576 |
| Guianan lowland moist forests | 0.956213 |
| Okhotsk-Manchurian taiga | 0.956468 |
| Southern Hudson Bay taiga | 0.956893 |
| Interior Yukon-Alaska alpine tundra | 0.957267 |
| Kaokoveld desert | 0.957367 |
| Canadian Low Arctic tundra | 0.957926 |
| Pantepui forests & shrublands | 0.95838 |
| Simpson desert | 0.958776 |
| Negro-Branco moist forests | 0.959608 |
| Red Sea coastal desert | 0.960212 |
| Cape York Peninsula tropical savanna | 0.962161 |
| Kalaallit Nunaat Arctic steppe | 0.962568 |
| Saharan Atlantic coastal desert | 0.962752 |
| Russian Bering tundra | 0.96279 |
| Guianan Highlands moist forests | 0.965348 |
| Kamchatka tundra | 0.966425 |
| Great Victoria desert | 0.966742 |
| Taimyr-Central Siberian tundra | 0.967773 |
| Western Australian Mulga shrublands | 0.969251 |
| Kalaallit Nunaat High Arctic tundra | 0.969928 |
| Russian Arctic desert | 0.97005 |
| Kamchatka taiga | 0.970495 |
| Japurá-Solimões-Negro moist forests | 0.971908 |
| Carnarvon xeric shrublands | 0.971991 |
| West Sahara desert | 0.972045 |
| East Siberian taiga | 0.972739 |
| Muskwa-Slave Lake taiga | 0.975868 |
| Trans-Baikal Bald Mountain tundra | 0.975884 |
| Solimões-Japurá moist forests | 0.975924 |
| Watson Highlands taiga | 0.97719 |
| South Sahara desert | 0.977261 |
| Canadian Middle Arctic Tundra | 0.97879 |
| Torngat Mountain tundra | 0.97932 |
| West Saharan montane xeric woodlands | 0.979877 |
| East Sahara Desert | 0.982581 |
| Cherskii-Kolyma mountain tundra | 0.982869 |
| Davis Highlands tundra | 0.983567 |
| Northeast Siberian taiga | 0.983774 |
| Namib Desert | 0.986127 |
| Juruá-Purus moist forests | 0.987731 |

| | |
|---|---|
| Canadian High Arctic tundra | 0.990033 |
| Tibesti-Jebel Uweinat montane xeric woodlands | 0.991717 |
| Ogilvie-MacKenzie alpine tundra | 0.992125 |
| Rock and Ice | 0.99417 |

**B)**

| **Ecosystem name** | **MEAN EII** |
|---|---|
| Warm Temperate Dry Settlement on Plains | 0.167071 |
| Tropical Moist Settlement on Plains | 0.185243 |
| Warm Temperate Moist Settlement on Plains | 0.194053 |
| Tropical Dry Settlement on Tablelands | 0.21042 |
| Sub Tropical Desert Settlement on Plains | 0.211817 |
| Warm Temperate Dry Settlement on Tablelands | 0.212276 |
| Tropical Dry Settlement on Hills | 0.216175 |
| Tropical Dry Settlement on Plains | 0.216739 |
| Warm Temperate Dry Settlement on Hills | 0.226508 |
| Sub Tropical Dry Settlement on Plains | 0.226882 |
| Sub Tropical Moist Settlement on Hills | 0.227386 |
| Tropical Moist Settlement on Tablelands | 0.232651 |
| Tropical Moist Settlement on Hills | 0.234444 |
| Warm Temperate Dry Settlement on Mountains | 0.241292 |
| Sub Tropical Moist Settlement on Plains | 0.241705 |
| Cool Temperate Dry Settlement on Tablelands | 0.241718 |
| Sub Tropical Dry Settlement on Tablelands | 0.242759 |
| Warm Temperate Moist Settlement on Hills | 0.247331 |
| Sub Tropical Moist Settlement on Tablelands | 0.252333 |
| Tropical Moist Settlement on Mountains | 0.25261 |
| Sub Tropical Dry Settlement on Hills | 0.253871 |
| Warm Temperate Moist Settlement on Tablelands | 0.25542 |
| Cool Temperate Moist Settlement on Plains | 0.255983 |
| Sub Tropical Moist Settlement on Mountains | 0.266251 |
| Sub Tropical Desert Settlement on Hills | 0.269048 |
| Cool Temperate Moist Settlement on Hills | 0.275133 |
| Warm Temperate Moist Settlement on Mountains | 0.27814 |
| Cool Temperate Moist Settlement on Tablelands | 0.278903 |
| Cool Temperate Dry Settlement on Hills | 0.284234 |
| Cool Temperate Dry Settlement on Plains | 0.28468 |
| Sub Tropical Dry Settlement on Mountains | 0.296984 |
| Polar Dry Settlement on Hills | 0.300584 |
| Warm Temperate Moist Cropland on Plains | 0.30413 |
| Sub Tropical Desert Settlement on Mountains | 0.30744 |
| Cool Temperate Dry Settlement on Mountains | 0.30968 |
| Tropical Dry Settlement on Mountains | 0.310982 |

| | |
|---|---|
| Warm Temperate Desert Settlement on Plains | 0.322074 |
| Warm Temperate Desert Forest on Tablelands | 0.333702 |
| Sub Tropical Desert Settlement on Tablelands | 0.33861 |
| Sub Tropical Desert Cropland on Plains | 0.342125 |
| Warm Temperate Dry Cropland on Plains | 0.348607 |
| Tropical Dry Cropland on Plains | 0.351783 |
| Cool Temperate Moist Settlement on Mountains | 0.362638 |
| Cool Temperate Moist Cropland on Plains | 0.368882 |
| Warm Temperate Moist Cropland on Hills | 0.370471 |
| Warm Temperate Desert Cropland on Plains | 0.373429 |
| Tropical Dry Cropland on Hills | 0.376593 |
| Polar Dry Settlement on Plains | 0.376927 |
| Warm Temperate Moist Sparsely or Non vegetated on Plains | 0.378447 |
| Cool Temperate Moist Cropland on Hills | 0.38484 |
| Tropical Moist Cropland on Plains | 0.390513 |
| Cool Temperate Moist Cropland on Tablelands | 0.393512 |
| Warm Temperate Desert Settlement on Tablelands | 0.398946 |
| Warm Temperate Dry Cropland on Tablelands | 0.409501 |
| Warm Temperate Dry Cropland on Hills | 0.410818 |
| Warm Temperate Dry Cropland on Mountains | 0.425929 |
| Tropical Dry Cropland on Tablelands | 0.42797 |
| Tropical Dry Cropland on Mountains | 0.432579 |
| Warm Temperate Moist Cropland on Mountains | 0.435387 |
| Warm Temperate Moist Sparsely or Non vegetated on Hills | 0.450611 |
| Sub Tropical Moist Cropland on Plains | 0.451557 |
| Cool Temperate Dry Cropland on Plains | 0.454088 |
| Sub Tropical Dry Cropland on Plains | 0.454965 |
| Warm Temperate Desert Cropland on Hills | 0.456435 |
| Warm Temperate Moist Cropland on Tablelands | 0.458033 |
| Tropical Moist Sparsely or Non vegetated on Mountains | 0.459456 |
| Cool Temperate Dry Cropland on Tablelands | 0.461623 |
| Cool Temperate Dry Cropland on Hills | 0.466254 |
| Cool Temperate Desert Settlement on Plains | 0.466486 |
| Warm Temperate Desert Cropland on Tablelands | 0.472613 |
| Tropical Desert Settlement on Plains | 0.476059 |
| Tropical Moist Cropland on Mountains | 0.476447 |
| Tropical Desert Settlement on Mountains | 0.489317 |
| Sub Tropical Desert Cropland on Mountains | 0.494134 |
| Cool Temperate Moist Grassland on Hills | 0.505686 |
| Sub Tropical Dry Cropland on Hills | 0.50665 |
| Sub Tropical Moist Cropland on Tablelands | 0.508087 |
| Warm Temperate Desert Settlement on Hills | 0.508747 |
| Sub Tropical Dry Cropland on Mountains | 0.509147 |
| Sub Tropical Moist Cropland on Hills | 0.512492 |

| | |
|---|---|
| Tropical Desert Settlement on Tablelands | 0.517291 |
| Cool Temperate Moist Grassland on Tablelands | 0.521347 |
| Warm Temperate Moist Shrubland on Plains | 0.52142 |
| Cool Temperate Dry Cropland on Mountains | 0.524122 |
| Cool Temperate Moist Cropland on Mountains | 0.524862 |
| Tropical Moist Cropland on Hills | 0.525558 |
| Warm Temperate Desert Settlement on Mountains | 0.525609 |
| Sub Tropical Moist Sparsely or Non vegetated on Plains | 0.525736 |
| Sub Tropical Dry Cropland on Tablelands | 0.527483 |
| Tropical Moist Sparsely or Non vegetated on Hills | 0.527901 |
| Tropical Desert Cropland on Mountains | 0.53012 |
| Sub Tropical Moist Cropland on Mountains | 0.530617 |
| Tropical Desert Cropland on Plains | 0.530619 |
| Sub Tropical Moist Sparsely or Non vegetated on Tablelands | 0.530809 |
| Tropical Moist Sparsely or Non vegetated on Tablelands | 0.533322 |
| Tropical Desert Settlement on Hills | 0.538876 |
| Tropical Moist Cropland on Tablelands | 0.547617 |
| Cool Temperate Moist Grassland on Plains | 0.561388 |
| Warm Temperate Moist Sparsely or Non vegetated on Mountains | 0.565719 |
| Warm Temperate Desert Forest on Mountains | 0.572217 |
| Warm Temperate Moist Grassland on Plains | 0.574176 |
| Boreal Moist Settlement on Hills | 0.577427 |
| Warm Temperate Desert Cropland on Mountains | 0.592336 |
| Cool Temperate Desert Cropland on Plains | 0.594187 |
| Warm Temperate Moist Grassland on Hills | 0.595914 |
| Warm Temperate Moist Sparsely or Non vegetated on Tablelands | 0.599411 |
| Polar Moist Settlement on Mountains | 0.600409 |
| Polar Dry Settlement on Mountains | 0.602286 |
| Warm Temperate Moist Shrubland on Hills | 0.604256 |
| Warm Temperate Dry Snow and Ice on Mountains | 0.60446 |
| Boreal Moist Settlement on Plains | 0.60809 |
| Sub Tropical Desert Cropland on Hills | 0.619873 |
| Warm Temperate Moist Forest on Plains | 0.621941 |
| Warm Temperate Moist Grassland on Tablelands | 0.62494 |
| Tropical Moist Sparsely or Non vegetated on Plains | 0.625468 |
| Sub Tropical Moist Grassland on Tablelands | 0.626821 |
| Warm Temperate Moist Grassland on Mountains | 0.629395 |
| Tropical Moist Grassland on Mountains | 0.632456 |
| Sub Tropical Desert Forest on Plains | 0.633162 |
| Boreal Moist Settlement on Tablelands | 0.635782 |
| Polar Moist Settlement on Hills | 0.636842 |
| Sub Tropical Moist Grassland on Mountains | 0.637599 |
| Sub Tropical Moist Sparsely or Non vegetated on Mountains | 0.64213 |

| | |
|---|---|
| Boreal Moist Cropland on Hills | 0.642161 |
| Warm Temperate Moist Forest on Hills | 0.642387 |
| Sub Tropical Moist Grassland on Hills | 0.645144 |
| Warm Temperate Moist Forest on Mountains | 0.648911 |
| Warm Temperate Moist Shrubland on Mountains | 0.650051 |
| Sub Tropical Desert Cropland on Tablelands | 0.651523 |
| Sub Tropical Moist Grassland on Plains | 0.661209 |
| Warm Temperate Dry Grassland on Plains | 0.662083 |
| Tropical Desert Cropland on Hills | 0.666574 |
| Boreal Dry Settlement on Hills | 0.67159 |
| Boreal Dry Settlement on Mountains | 0.671754 |
| Cool Temperate Moist Grassland on Mountains | 0.675159 |
| Warm Temperate Desert Forest on Plains | 0.678588 |
| Boreal Moist Cropland on Plains | 0.679436 |
| Warm Temperate Moist Forest on Tablelands | 0.681575 |
| Tropical Dry Forest on Mountains | 0.683828 |
| Warm Temperate Moist Shrubland on Tablelands | 0.685529 |
| Warm Temperate Dry Grassland on Hills | 0.685963 |
| Sub Tropical Moist Sparsely or Non vegetated on Hills | 0.687234 |
| Boreal Moist Cropland on Tablelands | 0.689472 |
| Cool Temperate Desert Cropland on Hills | 0.690253 |
| Boreal Moist Settlement on Mountains | 0.690457 |
| Cool Temperate Dry Grassland on Mountains | 0.699699 |
| Sub Tropical Desert Forest on Mountains | 0.702024 |
| Boreal Dry Settlement on Plains | 0.702329 |
| Sub Tropical Moist Shrubland on Plains | 0.703239 |
| Warm Temperate Dry Grassland on Mountains | 0.704307 |
| Boreal Dry Settlement on Tablelands | 0.705813 |
| Cool Temperate Dry Grassland on Tablelands | 0.71878 |
| Cool Temperate Moist Sparsely or Non vegetated on Hills | 0.719098 |
| Sub Tropical Moist Shrubland on Mountains | 0.719363 |
| Polar Desert Settlement on Mountains | 0.720035 |
| Warm Temperate Dry Grassland on Tablelands | 0.72045 |
| Warm Temperate Dry Forest on Mountains | 0.721033 |
| Tropical Dry Forest on Tablelands | 0.721259 |
| Tropical Moist Shrubland on Mountains | 0.721798 |
| Tropical Moist Grassland on Hills | 0.724465 |
| Sub Tropical Desert Forest on Hills | 0.726599 |
| Tropical Moist Grassland on Tablelands | 0.727881 |
| Cool Temperate Dry Grassland on Plains | 0.727903 |
| Cool Temperate Dry Grassland on Hills | 0.728388 |
| Sub Tropical Moist Shrubland on Hills | 0.730397 |
| Cool Temperate Dry Forest on Plains | 0.734921 |
| Tropical Dry Grassland on Plains | 0.737741 |

| | |
|---|---|
| Warm Temperate Dry Forest on Tablelands | 0.741291 |
| Cool Temperate Desert Cropland on Tablelands | 0.743346 |
| Sub Tropical Moist Forest on Mountains | 0.745738 |
| Sub Tropical Moist Shrubland on Tablelands | 0.747659 |
| Cool Temperate Moist Sparsely or Non vegetated on Plains | 0.748441 |
| Tropical Dry Grassland on Hills | 0.750426 |
| Sub Tropical Dry Forest on Mountains | 0.755379 |
| Cool Temperate Desert Settlement on Hills | 0.755557 |
| Boreal Dry Cropland on Hills | 0.757157 |
| Tropical Dry Forest on Hills | 0.757309 |
| Cool Temperate Dry Forest on Hills | 0.757745 |
| Tropical Dry Shrubland on Mountains | 0.759984 |
| Cool Temperate Moist Forest on Tablelands | 0.760115 |
| Tropical Dry Grassland on Mountains | 0.76292 |
| Sub Tropical Desert Grassland on Plains | 0.763327 |
| Warm Temperate Dry Sparsely or Non vegetated on Mountains | 0.763631 |
| Cool Temperate Moist Sparsely or Non vegetated on Tablelands | 0.765007 |
| Sub Tropical Dry Grassland on Mountains | 0.766657 |
| Tropical Dry Grassland on Tablelands | 0.767858 |
| Warm Temperate Desert Shrubland on Plains | 0.768166 |
| Warm Temperate Desert Grassland on Plains | 0.772286 |
| Cool Temperate Moist Sparsely or Non vegetated on Mountains | 0.772825 |
| Cool Temperate Moist Shrubland on Mountains | 0.772843 |
| Warm Temperate Dry Forest on Hills | 0.773858 |
| Cool Temperate Moist Forest on Mountains | 0.773863 |
| Tropical Moist Grassland on Plains | 0.774132 |
| Boreal Dry Cropland on Tablelands | 0.774297 |
| Polar Moist Cropland on Plains | 0.775849 |
| Tropical Dry Forest on Plains | 0.778424 |
| Cool Temperate Moist Forest on Plains | 0.778746 |
| Sub Tropical Moist Forest on Plains | 0.779977 |
| Boreal Dry Cropland on Plains | 0.782607 |
| Tropical Moist Forest on Mountains | 0.783649 |
| Polar Moist Cropland on Tablelands | 0.78698 |
| Tropical Desert Cropland on Tablelands | 0.787195 |
| Cool Temperate Moist Forest on Hills | 0.787974 |
| Sub Tropical Dry Shrubland on Mountains | 0.788559 |
| Tropical Moist Shrubland on Hills | 0.790018 |
| Tropical Dry Shrubland on Hills | 0.79003 |
| Tropical Moist Shrubland on Plains | 0.790239 |
| Warm Temperate Dry Sparsely or Non vegetated on Tablelands | 0.791187 |
| Tropical Dry Shrubland on Plains | 0.791595 |
| Tropical Desert Shrubland on Plains | 0.7987 |
| Warm Temperate Dry Shrubland on Mountains | 0.799102 |

| | |
|---|---|
| Sub Tropical Dry Forest on Plains | 0.800201 |
| Warm Temperate Desert Grassland on Mountains | 0.802236 |
| Sub Tropical Dry Shrubland on Tablelands | 0.80245 |
| Tropical Desert Forest on Mountains | 0.802578 |
| Tropical Dry Sparsely or Non vegetated on Mountains | 0.803334 |
| Warm Temperate Dry Sparsely or Non vegetated on Hills | 0.80767 |
| Sub Tropical Dry Sparsely or Non vegetated on Mountains | 0.808677 |
| Cool Temperate Moist Shrubland on Hills | 0.808901 |
| Sub Tropical Dry Shrubland on Hills | 0.812048 |
| Cool Temperate Desert Cropland on Mountains | 0.812224 |
| Warm Temperate Dry Shrubland on Tablelands | 0.812288 |
| Sub Tropical Moist Forest on Tablelands | 0.813513 |
| Cool Temperate Moist Shrubland on Tablelands | 0.815274 |
| Cool Temperate Moist Shrubland on Plains | 0.817367 |
| Sub Tropical Moist Forest on Hills | 0.819207 |
| Sub Tropical Dry Forest on Hills | 0.820605 |
| Polar Moist Forest on Mountains | 0.82292 |
| Sub Tropical Dry Grassland on Tablelands | 0.823504 |
| Warm Temperate Desert Forest on Hills | 0.824003 |
| Polar Moist Cropland on Mountains | 0.825371 |
| Cool Temperate Dry Forest on Mountains | 0.825415 |
| Tropical Moist Shrubland on Tablelands | 0.826038 |
| Warm Temperate Dry Shrubland on Hills | 0.828237 |
| Warm Temperate Dry Shrubland on Plains | 0.829213 |
| Warm Temperate Dry Forest on Plains | 0.831275 |
| Tropical Dry Shrubland on Tablelands | 0.83164 |
| Sub Tropical Dry Forest on Tablelands | 0.833665 |
| Cool Temperate Dry Shrubland on Hills | 0.834733 |
| Cool Temperate Dry Shrubland on Mountains | 0.836065 |
| Cool Temperate Dry Shrubland on Tablelands | 0.836542 |
| Cool Temperate Dry Shrubland on Plains | 0.836655 |
| Warm Temperate Dry Sparsely or Non vegetated on Plains | 0.837647 |
| Tropical Desert Shrubland on Mountains | 0.838258 |
| Sub Tropical Dry Shrubland on Plains | 0.839692 |
| Tropical Desert Forest on Plains | 0.841302 |
| Polar Dry Forest on Tablelands | 0.841847 |
| Polar Moist Grassland on Mountains | 0.841966 |
| Tropical Desert Grassland on Plains | 0.842221 |
| Warm Temperate Desert Grassland on Tablelands | 0.84327 |
| Cool Temperate Desert Settlement on Mountains | 0.843779 |
| Polar Dry Cropland on Tablelands | 0.843885 |
| Cool Temperate Dry Forest on Tablelands | 0.845329 |
| Cool Temperate Dry Sparsely or Non vegetated on Mountains | 0.845517 |
| Tropical Desert Grassland on Mountains | 0.850678 |

| | |
|---|---|
| Boreal Dry Cropland on Mountains | 0.851282 |
| Boreal Dry Grassland on Tablelands | 0.851895 |
| Boreal Dry Grassland on Hills | 0.855209 |
| Tropical Moist Forest on Tablelands | 0.857089 |
| Warm Temperate Desert Shrubland on Hills | 0.857178 |
| Polar Dry Forest on Mountains | 0.858031 |
| Polar Dry Cropland on Mountains | 0.860403 |
| Tropical Dry Sparsely or Non vegetated on Hills | 0.860591 |
| Tropical Dry Sparsely or Non vegetated on Tablelands | 0.862219 |
| Sub Tropical Dry Grassland on Hills | 0.863685 |
| Tropical Moist Forest on Hills | 0.865025 |
| Sub Tropical Desert Shrubland on Plains | 0.865344 |
| Boreal Dry Grassland on Mountains | 0.86543 |
| Polar Moist Grassland on Tablelands | 0.865516 |
| Sub Tropical Desert Shrubland on Hills | 0.865575 |
| Polar Moist Shrubland on Mountains | 0.866633 |
| Boreal Moist Cropland on Mountains | 0.866793 |
| Sub Tropical Dry Sparsely or Non vegetated on Tablelands | 0.869148 |
| Cool Temperate Dry Sparsely or Non vegetated on Plains | 0.869462 |
| Cool Temperate Desert Grassland on Plains | 0.870283 |
| Cool Temperate Desert Grassland on Mountains | 0.870632 |
| Cool Temperate Dry Snow and Ice on Mountains | 0.870839 |
| Cool Temperate Moist Snow and Ice on Hills | 0.871294 |
| Boreal Dry Sparsely or Non vegetated on Hills | 0.871635 |
| Polar Moist Cropland on Hills | 0.871767 |
| Polar Moist Forest on Tablelands | 0.871887 |
| Tropical Desert Grassland on Hills | 0.873225 |
| Tropical Desert Forest on Hills | 0.873536 |
| Boreal Dry Sparsely or Non vegetated on Mountains | 0.874601 |
| Polar Moist Forest on Hills | 0.875962 |
| Warm Temperate Desert Grassland on Hills | 0.877283 |
| Tropical Dry Sparsely or Non vegetated on Plains | 0.877466 |
| Cool Temperate Desert Grassland on Tablelands | 0.878446 |
| Polar Dry Cropland on Hills | 0.879479 |
| Sub Tropical Desert Shrubland on Mountains | 0.879866 |
| Boreal Dry Sparsely or Non vegetated on Tablelands | 0.8799 |
| Tropical Desert Shrubland on Hills | 0.880141 |
| Cool Temperate Dry Sparsely or Non vegetated on Tablelands | 0.881001 |
| Tropical Desert Grassland on Tablelands | 0.882121 |
| Cool Temperate Desert Forest on Mountains | 0.883043 |
| Warm Temperate Desert Shrubland on Mountains | 0.88335 |
| Tropical Moist Forest on Plains | 0.885684 |
| Boreal Dry Grassland on Plains | 0.886898 |
| Polar Moist Shrubland on Tablelands | 0.886918 |

| | |
|---|---|
| Cool Temperate Moist Snow and Ice on Mountains | 0.886984 |
| Warm Temperate Moist Snow and Ice on Mountains | 0.887759 |
| Cool Temperate Dry Sparsely or Non vegetated on Hills | 0.889487 |
| Polar Dry Cropland on Plains | 0.890423 |
| Sub Tropical Dry Sparsely or Non vegetated on Hills | 0.892388 |
| Polar Moist Grassland on Hills | 0.892456 |
| Polar Dry Grassland on Mountains | 0.89269 |
| Boreal Desert Grassland on Mountains | 0.89424 |
| Sub Tropical Dry Grassland on Plains | 0.894429 |
| Tropical Desert Shrubland on Tablelands | 0.89495 |
| Polar Dry Grassland on Hills | 0.89642 |
| Tropical Desert Sparsely or Non vegetated on Mountains | 0.899439 |
| Polar Dry Grassland on Tablelands | 0.900172 |
| Tropical Desert Forest on Tablelands | 0.900708 |
| Polar Dry Shrubland on Mountains | 0.901042 |
| Polar Desert Settlement on Tablelands | 0.902378 |
| Boreal Desert Cropland on Mountains | 0.903876 |
| Cool Temperate Desert Grassland on Hills | 0.904456 |
| Cool Temperate Moist Snow and Ice on Tablelands | 0.904507 |
| Polar Dry Settlement on Tablelands | 0.904686 |
| Boreal Moist Grassland on Hills | 0.906176 |
| Boreal Moist Grassland on Plains | 0.906489 |
| Warm Temperate Desert Sparsely or Non vegetated on Mountains | 0.906947 |
| Polar Moist Shrubland on Hills | 0.908986 |
| Polar Dry Grassland on Plains | 0.909703 |
| Sub Tropical Desert Grassland on Hills | 0.912599 |
| Boreal Moist Grassland on Mountains | 0.912624 |
| Sub Tropical Desert Grassland on Mountains | 0.914888 |
| Sub Tropical Dry Sparsely or Non vegetated on Plains | 0.917027 |
| Polar Dry Shrubland on Hills | 0.918365 |
| Boreal Moist Grassland on Tablelands | 0.918718 |
| Sub Tropical Desert Shrubland on Tablelands | 0.920351 |
| Polar Dry Shrubland on Tablelands | 0.922165 |
| Cool Temperate Desert Snow and Ice on Mountains | 0.922208 |
| Cool Temperate Desert Shrubland on Tablelands | 0.923556 |
| Warm Temperate Desert Shrubland on Tablelands | 0.926497 |
| Polar Moist Grassland on Plains | 0.928598 |
| Boreal Moist Shrubland on Plains | 0.929546 |
| Polar Desert Cropland on Mountains | 0.929952 |
| Polar Desert Sparsely or Non vegetated on Tablelands | 0.931233 |
| Cool Temperate Desert Forest on Plains | 0.931525 |
| Boreal Dry Shrubland on Mountains | 0.932763 |
| Polar Desert Grassland on Mountains | 0.933177 |
| Polar Dry Forest on Hills | 0.933991 |

| | |
|---|---|
| Sub Tropical Desert Grassland on Tablelands | 0.934061 |
| Sub Tropical Desert Sparsely or Non vegetated on Mountains | 0.935362 |
| Boreal Dry Sparsely or Non vegetated on Plains | 0.93674 |
| Cool Temperate Desert Shrubland on Plains | 0.93725 |
| Polar Moist Settlement on Plains | 0.93739 |
| Cool Temperate Desert Shrubland on Mountains | 0.938913 |
| Warm Temperate Desert Sparsely or Non vegetated on Plains | 0.939832 |
| Polar Desert Snow and Ice on Mountains | 0.94064 |
| Polar Dry Sparsely or Non vegetated on Mountains | 0.940859 |
| Boreal Dry Snow and Ice on Mountains | 0.941007 |
| Polar Dry Shrubland on Plains | 0.941044 |
| Warm Temperate Desert Sparsely or Non vegetated on Tablelands | 0.943122 |
| Boreal Dry Shrubland on Hills | 0.945329 |
| Boreal Moist Shrubland on Hills | 0.945988 |
| Tropical Desert Sparsely or Non vegetated on Hills | 0.946151 |
| Boreal Dry Shrubland on Plains | 0.947198 |
| Boreal Moist Sparsely or Non vegetated on Mountains | 0.947568 |
| Polar Desert Sparsely or Non vegetated on Mountains | 0.949041 |
| Boreal Moist Shrubland on Tablelands | 0.949212 |
| Tropical Desert Sparsely or Non vegetated on Tablelands | 0.949466 |
| Cool Temperate Desert Sparsely or Non vegetated on Mountains | 0.950219 |
| Boreal Moist Forest on Plains | 0.953081 |
| Tropical Desert Sparsely or Non vegetated on Plains | 0.95332 |
| Polar Moist Sparsely or Non vegetated on Mountains | 0.955021 |
| Boreal Dry Forest on Mountains | 0.955224 |
| Boreal Moist Shrubland on Mountains | 0.956577 |
| Polar Moist Shrubland on Plains | 0.956801 |
| Boreal Moist Forest on Hills | 0.95765 |
| Boreal Moist Sparsely or Non vegetated on Plains | 0.958497 |
| Sub Tropical Desert Sparsely or Non vegetated on Tablelands | 0.958662 |
| Boreal Dry Shrubland on Tablelands | 0.958779 |
| Boreal Moist Sparsely or Non vegetated on Tablelands | 0.95929 |
| Boreal Moist Sparsely or Non vegetated on Hills | 0.959379 |
| Polar Moist Forest on Plains | 0.960261 |
| Warm Temperate Desert Sparsely or Non vegetated on Hills | 0.960682 |
| Sub Tropical Desert Sparsely or Non vegetated on Hills | 0.960692 |
| Boreal Desert Sparsely or Non vegetated on Plains | 0.961059 |
| Polar Dry Forest on Plains | 0.961186 |
| Polar Desert Shrubland on Mountains | 0.961528 |
| Sub Tropical Desert Sparsely or Non vegetated on Plains | 0.961755 |
| Polar Desert Sparsely or Non vegetated on Plains | 0.962127 |
| Polar Dry Snow and Ice on Mountains | 0.962245 |
| Cool Temperate Desert Sparsely or Non vegetated on Plains | 0.962928 |

| | |
|---|---|
| Boreal Dry Forest on Tablelands | 0.964156 |
| Boreal Moist Forest on Mountains | 0.966071 |
| Boreal Desert Sparsely or Non vegetated on Mountains | 0.966188 |
| Boreal Desert Grassland on Plains | 0.966292 |
| Polar Moist Snow and Ice on Mountains | 0.966727 |
| Boreal Dry Forest on Hills | 0.966966 |
| Polar Desert Grassland on Plains | 0.967258 |
| Boreal Moist Forest on Tablelands | 0.968446 |
| Boreal Dry Forest on Plains | 0.969792 |
| Polar Dry Sparsely or Non vegetated on Plains | 0.97028 |
| Polar Desert Forest on Mountains | 0.97097 |
| Polar Moist Sparsely or Non vegetated on Plains | 0.974636 |
| Boreal Moist Snow and Ice on Mountains | 0.974814 |
| Cool Temperate Desert Sparsely or Non vegetated on Tablelands | 0.974995 |
| Cool Temperate Desert Shrubland on Hills | 0.976623 |
| Polar Desert Shrubland on Plains | 0.977507 |
| Polar Dry Sparsely or Non vegetated on Hills | 0.977726 |
| Polar Dry Sparsely or Non vegetated on Tablelands | 0.978717 |
| Cool Temperate Desert Sparsely or Non vegetated on Hills | 0.979301 |
| Polar Desert Shrubland on Tablelands | 0.98174 |
| Sub Tropical Desert Forest on Tablelands | 0.982381 |
| Polar Moist Sparsely or Non vegetated on Hills | 0.983771 |
| Polar Desert Cropland on Plains | 0.98522 |
| Polar Dry Snow and Ice on Plains | 0.985298 |
| Polar Desert Forest on Plains | 0.986246 |
| Polar Moist Sparsely or Non vegetated on Tablelands | 0.988936 |
| Boreal Desert Sparsely or Non vegetated on Tablelands | 0.99262 |
| Polar Dry Snow and Ice on Hills | 0.992968 |
| Cool Temperate Moist Snow and Ice on Plains | 0.99543 |
| Polar Dry Snow and Ice on Tablelands | 0.995641 |
| Polar Moist Snow and Ice on Hills | 0.998898 |
| Polar Moist Snow and Ice on Tablelands | 0.999125 |
| Boreal Moist Snow and Ice on Tablelands | 0.999142 |
| Polar Moist Snow and Ice on Plains | 0.9999 |
| Boreal Moist Snow and Ice on Plains | 0.999956 |
| Boreal Moist Snow and Ice on Hills | 0.999998 |

Disaggregation of components of EII

Compositional integrity estimates show that wide expanses of high-latitude regions have retained high compositional integrity (Figure S1A). The same is true for the desert and dryland regions of Africa and to a lesser extent Australia. However, species richness and regional endemism values are much lower in these regions when compared with the other broadly intact regions of the Amazon rainforest, parts of the Congo rainforest, and parts of the Indomalayan rainforest (especially on

Borneo and Papua New Guinea) (Burgess *et al.*, 2006; Fa & Funk, 2007). Notable regions of the world with degraded compositional integrity include India and China, Europe, eastern USA and Canada and high human population density regions of Africa and eastern South America.

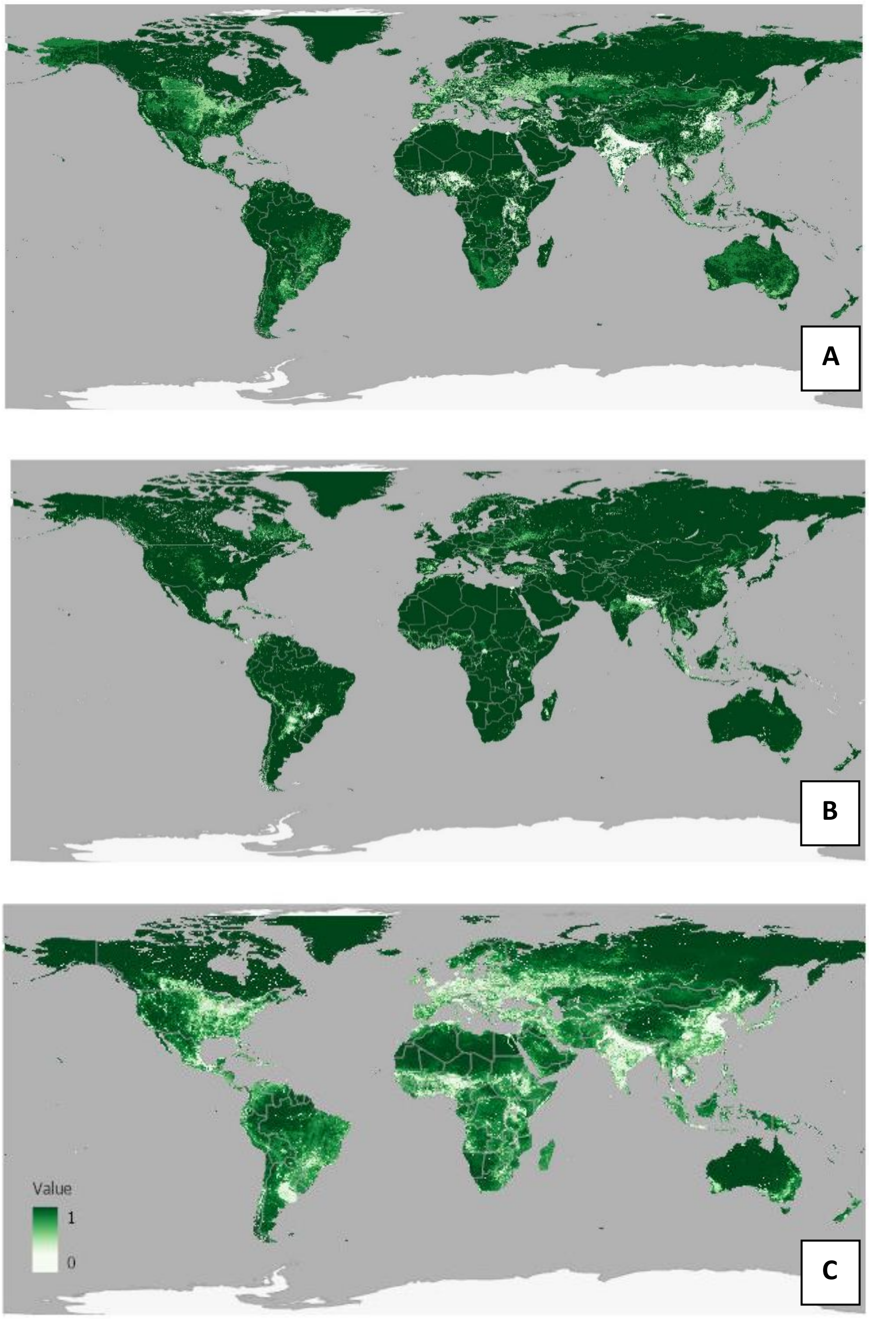


Figure S1A-C. Global maps of components of ecosystem integrity index: A) composition, B) functioning, and C) structure. Areas with high levels of integrity for each component part are shown in dark green; those with highly degraded integrity are shown in white.

Degradation to functional integrity (Figure S1B) is less widespread than degradation to compositional (Figure S1A) or structural integrity (Figure S1C). Highly degraded areas are often found where agriculture has replaced savanna, grasslands or scrub, for instance, those in the central agricultural regions of South America and Europe, in the Sahelian and Guinea Savanna region of Africa, and Central and north-eastern Asia and North America.

Structural integrity (Figure S1C) shows somewhat similar spatial patterns to compositional integrity; however, lower values of structural integrity are observed in areas of widespread intensive cropland or high human population density, for instance in throughout much of Europe, India, China and central-western North America.

Sensitivity analysis – comparison of EII layer with other global layers

Figure S2 shows the congruence between layers of anthropogenic impacts and EII by biome. There are few areas where of the world where all 3 comparison pressure layers agree that human pressures are low (shown in dark purple in plots); however, areas of agreement tended to be found in areas of high ecosystem integrity (values >0.8).  Areas where one or two of the layers showed low anthropogenic pressures still tended to correlate with high values of EII (values >0.7). Biomes showed differing patterns with clear bimodal alignment in tropical dry broadleaf forests, and, to a lesser extent, in temperate broadleaf forests, temperate grasslands and Mediterranean forests, woodland and scrub.

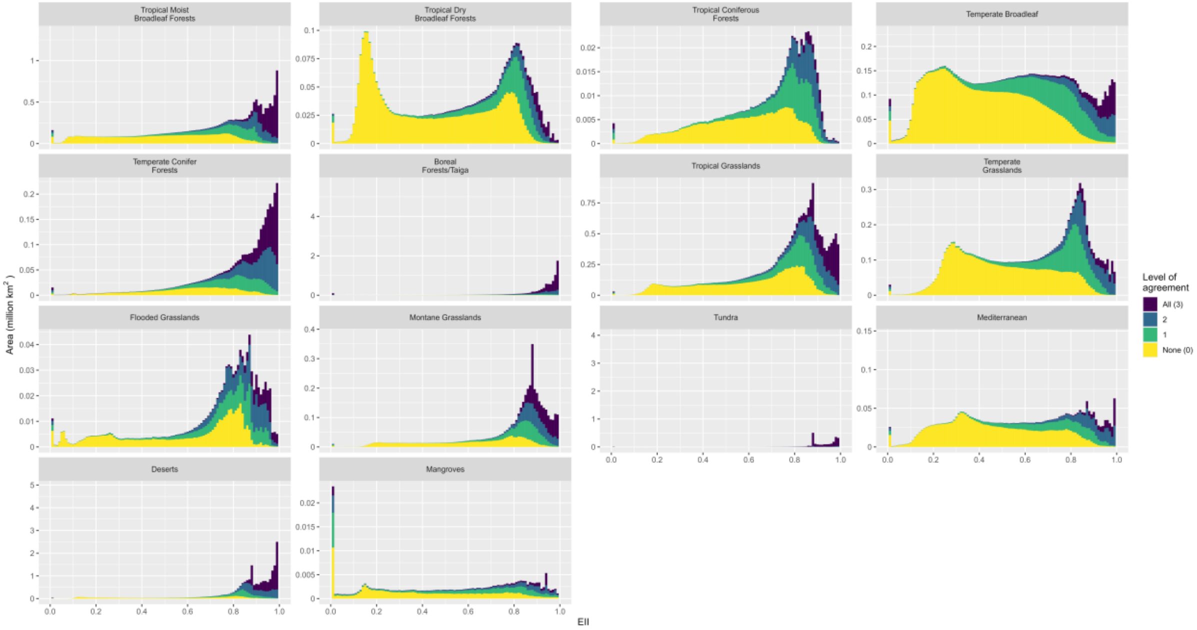


Figure S2. Comparison of EII values with A) Human Footprint Index (Watson *et al.,* 2020), B) natural and modified habitat (Gosling *et al.,* 2020), C) low impact areas (Jacobson *et al.*, 2019). Colours identify how many of the three comparison layers classify an area as experiencing low anthropogenic impact.

When comparing EII to the natural and modified habitat layer (Gosling *et al.,* 2020) expected patterns are observed with EII values >0.8 correlating with areas identified as 'likely natural', those areas identified as 'potentially natural' and 'potentially modified' tending to correlate with mid to high EII values, and areas identified as 'likely modified' tending to correlate with lower EII values (Figure S3).

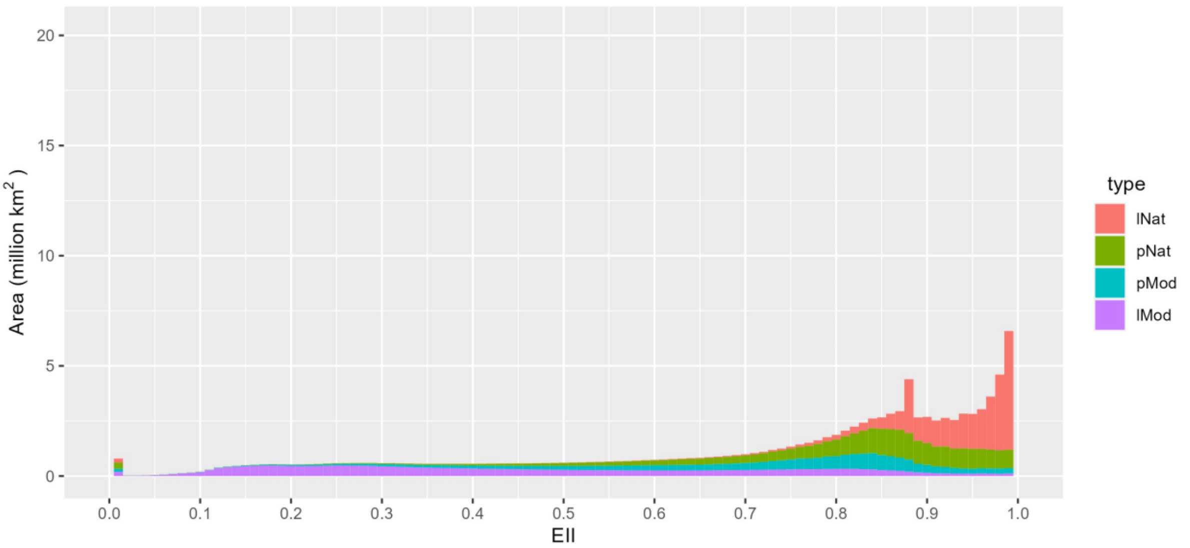


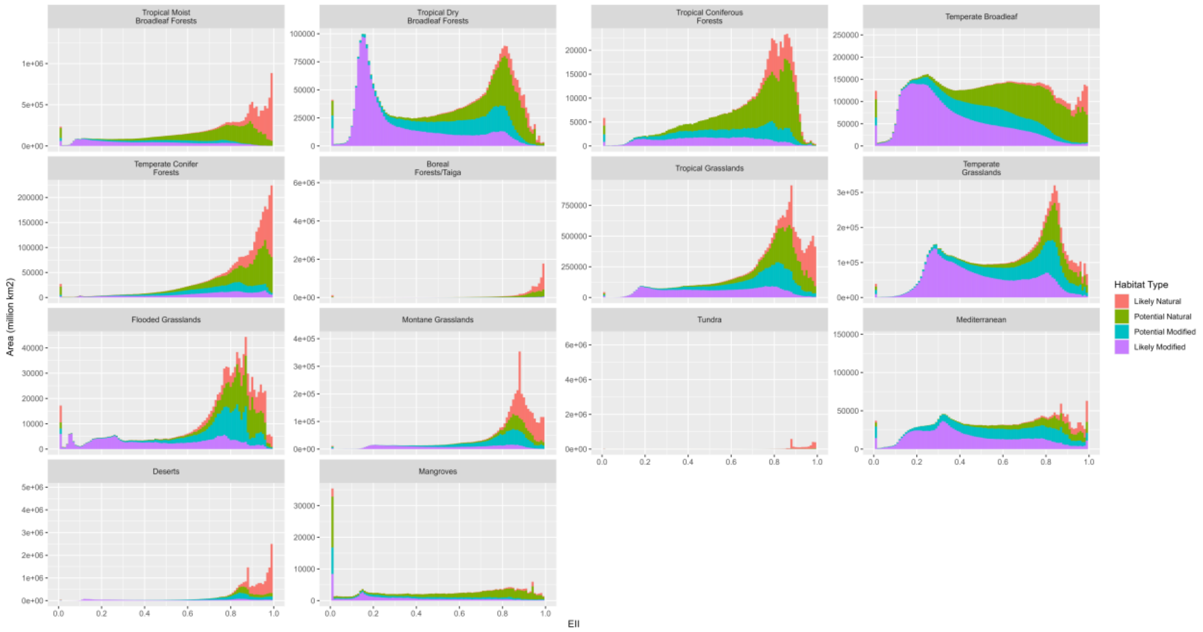


Figure S3. Comparison of EII values with the Natural and Modified Habitat layer developed by Gosling *et al.* (2020). Colours identify areas classified by Gosling *et al.* (2020) as ‘likely natural’, ‘potentially natural’, ‘potentially modified’ and ‘likely modified’.
